# METTL1 gene dosage and catalytic activity define a targetable vulnerability in pediatric cancers

**DOI:** 10.64898/2026.10.20.760582

**Authors:** Pei Y. Liu, Steven He, Olesya Panasenko, Fedor Bezrukov, Alice Salib, Priyank Rawat, Wenxin Hu, Chelsea Mayoh, Marie Erasmus-Wong, Zara Barger, Emmy Fleuren, Greg M. Arndt, Ian Street, Mohamed Fareh, Antoine de Weck, Paul G. Ekert

## Abstract

METTL1 is responsible for catalyzing N7-methylguanosine (m^7^G) methylation of tRNAs. METTL1-dependent oncogenic mechanisms include both its methyltransferase catalytic activity and, in specific instances, non-catalytic interactions that maintain aminoacylated tRNA levels in malignant cells. We show that *METTL1* copy number gain and elevated expression are a feature of pediatric tumors that are *METTL1* dependent, using Cas13-mediated RNA to reduce the threshold of METTL1 mRNA and protein, which disrupts both catalytic and non-catalytic functions. In cell lines with *METTL1* gene amplification, specific degradation of *METTL1* mRNA impaired colony formation, triggered cell death, and decreased the translation efficiency of a subset of genes enriched for codons decoded m^7^G-modified tRNAs. METTL1 inhibition was accompanied by downregulation of c-MYC, n-MYC and MCL-1 protein expression. METTL1 silencing synergizes with drugs targeting anti-apoptotic members of the BCL2 family, notably BCL-xL. Importantly, the catalytic activity of METTL1 was required to rescue cells from the deleterious consequences of METTL1 silencing. We conclude that inhibiting the catalytic activity of METTL1 has potential therapeutic efficacy, particularly in tumors with *METTL1* amplification. Synergistic combinations with BH3-mimetic drugs may be a promising therapeutic combination.

**Significance:** METTL1 amplification defines a subset of pediatric tumors. METTL1 inhibition using CRISPR-Cas13d induces cell death and synergizes with a BCL2/BCL-xL inhibitor, revealing a targetable dependency and a potential treatment combination.

## INTRODUCTION

Therapies targeting epigenetic regulators in cancer have primarily focused on DNA and histone modifications (1). However, enzymes modifying RNA are emerging therapeutic targets (2). More than 170 chemical modifications of RNAs have been identified (3), of which RNA methylation is one of the most common (4). RNA methyltransferases play crucial roles in transcription, RNA stability, translation, splicing and epigenetic regulation (5). The commonest RNA modifications in eukaryotes include N6-methyladenosine (m^6^A) - methylation of the nitrogen atom at the sixth position of adenosine (6), and N7-methylguanosine (m^7^G) - methylation of the nitrogen atom at the seventh position of guanosine (7).

The most studied m^6^A writer is METTL3, which exists in a complex with METTL14 and Wilms Tumor 1-associating protein (WTAP) (8). Elevated METTL3 expression is associated with resistance to radiotherapy in glioma (9) and immunotherapy resistance in liver cancer (10). Three other m^6^A methyltransferases are known. METTL16 catalyzes m^6^A methylation on U6 small nucleolar RNA to regulate RNA splicing (11) and has apparent driver roles in a range of cancers (12). METTL5 complexes with TRMT112 and is responsible for catalyzing m^6^A methylation on 18S rRNA (13). Upregulation of METTL5 promotes gastric cancer and breast cancer cell growth (14). Zinc finger CCHC domain-containing protein 4 (ZCCHC4) primarily methylates 28S rRNA (15). High ZCCHC4 expression is associated with chemoresistance in hepatocellular carcinoma patients by interrupting DNA damage-mediated cell death (16).

Three m^7^G methyltransferases are known. BUD23 facilitates m^7^G methylation on 18S rRNA at nucleotide G1639 by interacting with methyltransferase activator subunit 11–2 (TRMT112) (17). RNA guanine-7 methyltransferase (RNMT) catalyzes m^7^G methylation on the 5’ cap of mRNAs by cooperating with RNMT-activating miniprotein (RAM) (18). METTL1 mediates internal m^7^G methylation on tRNAs, mRNAs and miRNAs by interacting with tryptophan-aspartic (WD) repeat domain 4 (WDR4) (19). METTL1 overexpression increases tRNA stability and results in codon-biased translation of associated mRNAs and tumorigenesis, of which m^7^G methylation of Arg-TCT tRNA may be particularly important to its oncogenic function (19). Elevated METTL1 expression correlates with poor outcomes in sarcoma, glioblastoma multiforme and hepatocellular carcinoma (19). Overexpression, silencing and mutagenesis studies have linked METTL1 oncogenic function to glioma (20), lung cancer (21), intrahepatic cholangiocarcinoma (22), colorectal cancer (23), esophageal cancer (24) and liposarcoma (25). In most cases, with the notable exception of liposarcoma, the methyltransferase catalytic activity is mechanistically important. *METTL1* gene amplification is recognized in up to 20% of soft tissue sarcomas (26), glioblastoma (27) and rhabdomyosarcoma (28). Preclinical studies of a METTL1 inhibitory small molecule have demonstrated some efficacy, but have not yet proceeded to clinical trial (29).

Clustered Regularly Interspaced Short Palindromic Repeats (CRISPR) technology has been exploited for precise RNA editing and silencing (30). Cas13b (31) and Cas13d (32) are type VI CRISPR-Cas systems that target single-stranded RNAs for degradation via a programmable CRISPR RNA (crRNA) guide. The Cas13b and Cas13d crRNAs require 30bp and 23bp base-pairing respectively with the target mRNA to activate their nuclease activity, which offers significant specificity (33). Interestingly, various Cas13 systems can tolerate limited mismatches between the guide and the target RNA, enabling resilience against mutational escape that often drive resistance to anti-cancer or antiviral small molecule drugs. Unconventional design of Cas13d crRNA harboring synthetic mismatches has been used to specifically silence oncogenic mRNA transcribed from genes bearing point mutations such as KRAS G12, NRAS G12D and BRAF V600E without targeting non-mutated transcripts (34). We set out to interrogate METTL1 gene dosage across different cancer types with or without METTL1 amplification using RNA-targeting with CRISPR-Cas13 to dial METTL1 protein in amplified cells down to the levels seen in their non-amplified counterparts, and examined the consequences.

Here we show that *METTL1* is amplified and overexpressed in a subset of high-grade glioma, neuroblastoma, osteosarcoma and rhabdomyosarcoma tumors from pediatric cancer patients. We explored the potential of METTL1 as a therapeutic target in pediatric cancers using targeted Cas13d-driven mRNA degradation. METTL1 is a selective dependency gene in pediatric cancers, notably in selected neuroblastoma (NBL), osteosarcoma and rhabdomyosarcoma (RMS) cell lines with *METTL1* gene amplification. Cas13d-dependent silencing of *METTL1* mRNA was highly specific and reliably reduced METTL1 protein expression. In dependent cell lines, METTL1 silencing triggered cell death, diminished colony formation, and altered translation of a subset of genes in a codon-dependent manner. This dependency required METTL1 catalytic activity. Guided by an analysis of the transcriptional consequences of METTL1 silencing, we observed potential synergy between METTL1 inhibition and drugs acting on the intrinsic apoptosis pathway.

## MATERIALS AND METHODS

### Cell culture

BE(2)-C (RRID:CVCL_0529) neuroblastoma, RD rhabdomyosarcoma and HEK293T (RRID:CVCL_0063) cells were cultured in DMEM media supplemented with 10% fetal calf serum (FCS). TN-2 neuroblastoma (RRID:CVCL_5602) and Rh30 (RRID:CVCL_004) rhabdomyosarcoma cells were cultured in RPMI-1640 supplemented with 10% FCS. SJSA-1 (RRID:CVCL_1697) osteosarcoma cells were cultured in high glucose RPMI-1640 (Catalog number 30-2001, ATCC) media supplemented with 10% FCS. ZCCS1476 patient-derived osteosarcoma cells (OZ-SBT-1476) were cultured in IMDM (Catalog number 12440053, Gibco) media supplemented with 10% FCS, 1% Insulin-Transferrin-Selenium (Catalog number 41400045, Gibco) and 0.1% Rock inhibitor (Catalog number S6390, Sapphire). Tumor Stem Medium Base (TSM-B) was prepared from equal parts of neurobasal-A (Catalog number 10888022, Invitrogen) and DMEM/F-12 (Catalog number 11330032, Invitrogen), supplemented with 10 mM HEPES (Catalog number 15630080, Invitrogen), 1 mM sodium pyruvate (Catalog number 11360070, Invitrogen), 0.1 mM MEM non-essential amino acids (Catalog number 11140050, Invitrogen), 1× glutaMAX-I (Catalog number 35050061, Invitrogen) and 1× antibiotic-antimycotic (Catalog number 15240096, Invitrogen). AUS-DIPG-017 patient-derived diffuse intrinsic pontine glioma (DIPG) cells were cultured in TSM-B further supplemented (prepared fresh each time) with recombinant human EGF (20 ng/mL) (Catalog number AF-100-15-500 µg, PeproTech), FGF-basic (20 ng/mL) (Catalog number AF-100-18B-500 µg, PeproTech), PDGF-AA (10 ng/mL) (Catalog number AF-100-13A-100 µg, PeproTech), PDGF-BB (10 ng/mL) (Catalog number AF-100-14B-100 µg, PeproTech), heparin (2 µg/mL) (Catalog number 07980, StemCell Technologies) and 1× B-27 minus vitamin A (Catalog number 12587-010, Invitrogen). BE(2)-C cell line was a gift from Barbara Spengler (Fordham University, NY). TN-2 cell line was provided by the Riken BRC through the National BioResource Project of the MEXT, Japan. SJSA-1 cell line was purchased from ATCC. OZ-SBT-1476 cell line was generated by Emmy Fleuren’s group at Children’s Cancer Institute. AUS-DIPG-017 cell line was established by the Brain Cancer Team at Children’s Cancer Institute (35). RD and Rh30 cell lines were provided by Peter Houghton (Greehey Children’s Cancer Research Institute, The University of Texas at San Antonio, USA). All parental cell lines were authenticated in March 2025 or August 2026 by small tandem repeat (STR) profiling at CellBank Australia or Garvan Institute of Medical Research.

### Establishment of stable DOX-inducible CRISPR Cas13d effector and METTL1 CRISPR Cas13d guides in cell lines

The pLentiRNACRISPR_007-TetO-NLS-RfxCas13d-NLS-WPRE-EFS-rtTA3-2A-Blast (RRID:Addgene_138149) and pLentiRNAGuide_001-hU6-RfxCas13d-DR1-BsmBI-EFS-Puro-WPRE (RRID:Addgene_138150) constructs were obtained from Addgene. METTL1 guides were designed using the tool (https://cas13design.nygenome.org/) developed by Wessels et al (36). Forward and reverse negative control or METTL1 oligonucleotides were cloned into the pLentiRNAGuide_001-hU6-RfxCas13d-DR1-BsmBI-EFS-Puro-WPRE vector. HEK293T cells were transfected with pLentiRNACRISPR_007-TetO-NLS-RfxCas13d-NLS-WPRE-EFS-rtTA3-2A-Blast construct together with the pCMV-VSV-G envelope and pCMV-dR8.2 packaging plasmids. Virus-containing medium was collected and used to transduce BE(2)-C, TN-2, RD, Rh30, AUS-DIPG-017, SJSA-1 and OZ-SBT-1476 cells in the presence of polybrene (8 µg/mL). Transduced cells were selected with blasticidin S at concentrations and durations optimized for each cell line: BE(2)-C, 20 µg/mL for 7 days; RD, 20 µg/mL for 13 days; TN-2, SJSA-1 and Rh30, 10 µg/mL for 9 days; AUS-DIPG-017, 15 µg/mL for 4 days; OZ-SBT-1476, 2.5 µg/mL for 7 days. Subsequently, HEK293T cells were transfected with pLentiRNAGuide_001-hU6-RfxCas13d-DR1-BsmBI-EFS-Puro-WPRE in conjunction with the pCMV-VSV-G (RRID:Addgene_8454) envelope and pCMV-dR8.2 (RRID:Addgene_8455) packaging plasmids. Virus-containing medium was collected and used to transduce BE(2)-C, TN-2, RD, Rh30, OZ-SBT-1476, SJSA-1 and AUS-DIPG-017 cells expressing DOX-inducible Cas13d in the presence of polybrene. Transduced cells were then selected with puromycin for 3-7 days at 5 µg/mL [BE(2)-C, TN-2, RD, Rh30 and SJSA-1], 2.5 µg/mL (AUS-DIPG-017) or 1.5 µg/mL (OZ-SBT-1476). The non-targeting control guide sequence is CCGCACTGGTCTAAGCCGAAGAA, and the eight METTL1 target sequences are g1_ATGGCGGACCACACGCTGCGCTA, g2_TCGCTCCACTCACTCAAAATCAG, g3_ACGCTGCGCTACCCTGTGAAGCC, g4_CTGCGCTACCCTGTGAAGCCAGA, g5_CAAGTGGCGAATCATCAGTCCCA, g6_CTAGCAGAATATGCCTACGTGCT, g7_GCGCTAGTGAGGAGTGGGCAGAG and g8_ACACGCTGCGCTAGTGAGGAGTG.

### Quantitative real-time reverse transcription polymerase chain reaction (RT-PCR)

RT-PCR was carried out using Power SYBR Green Master Mix (ThermoFisher Scientific) with primers targeting specific genes on QuanStudio^TM^. Relative gene expression was calculated using the double delta Ct (_△△_Ct) method (37), after normalizing to the reference gene actin or GAPDH. The primers were synthesized by Merck Life Science and the sequences are: 5’-GCTGCGCTACCCTGTGAAG-3’ (forward) and 5’-TTGAGTGAGTGGAGCGAAGAAC-3’ (reverse) for METTL1 targeting amplicon 1; 5’-GCACAAGTGGCGAATCATCA-3’ (forward) and 5’-CAGCCCCCCAACTCTTAGC-3’ (reverse) for METTL1 targeting amplicon 2; 5’-GCACCGTCAAGGCTGAGAAC-3’ (forward) and 5’-GGATCTCGCTCCTGGAAGATG-3’ (reverse) for GAPDH; 5’-AGGCCAACCGCGAGAAG-3’ (forward) and 5’-ACAGCCTGGATAGCAACGTACA-3’ (reverse) for Actin.

### Western blot

DOX-inducible Cas13d cells with constitutive non-targeting control or METTL1 guide 2, 4 or 5 were cultured with DMSO control or DOX [1 µg/mL for Rh30, SJSA-1, OZ-SBT-1476 and AUS-DIPG-017, and 2 µg/mL for TN-2, RD and BE(2)-C] for the indicated time. Cell pellets were collected by trypsinization or scraping and centrifugation, then lysed in RIPA lysis buffer (1% NP-40, 150 mM NaCl, 0.1% SDS, 50 mM Tris-Cl, pH 7.5, and 0.5% sodium deoxycholate) with protease inhibitors (Roche, Penzberg, Germany) and phosphatase inhibitors (Sigma, St Louis, MO). Cell debris was removed by centrifugation, and protein concentration was measured using the Pierce™ BCA Protein Assay Kit (Pierce, Rockford, IL). Subsequently, 20-40 µg of protein was denatured, loaded into a well of a 4-20% precast polyacrylamide gel, and ran at 100 V for 90 minutes. Protein was transferred to a nitrocellulose membrane (Cytiva, Marlborough, MA), and the membrane was blocked with 10% skim milk dissolved in phosphate-buffered saline (PBS).

The membrane was then incubated with primary antibody overnight at 4°C, washed with PBST, probed with HRP-conjugated secondary antibody for 1.5 hours, washed with PBST, and imaged using a ChemiDoc Imaging System (Bio-Rad, Hercules, CA). The primary antibody conditions were: rabbit anti-METTL1 antibody (1:1000) (Proteintech, catalog number 14994-1-AP, RRID:AB_2142013), mouse anti-HA antibody (1:500) (Santa Cruz Biotechnology, catalog number sc-7392, RRID:AB_627809, ), rabbit anti-cleaved caspase-3 antibody (1:100) (Cell Signaling Technology, catalog number 9664, , RRID:AB_2070042), mouse anti-N-MYC antibody (1:1000) (Santa Cruz Biotechnology, catalog number sc-53993, RRID:AB_831602), rabbit anti-BCL2 antibody (1:50) (Invitrogen, catalog number MA5-11757, RRID:AB_10978135, ), rabbit anti-MCL-1 antibody (1:2000) (Rockland, catalog number 600-401-394, RRID:AB_2266446, ), rabbit anti-BCL-xL antibody (1:2000) (Invitrogen, catalog number MA5-15142, RRID: AB_10979212), rabbit anti-c-MYC antibody (Cell Signaling Technology, catalog number 13987S, RRID:AB_2631168), mouse anti-GAPDH antibody (1:5000) (Santa Cruz Biotechnology, catalog number sc-365062, RRID:AB_10847862,), or mouse anti-actin antibody (1:5000) (Sigma-Aldrich, catalog number A2228, RRID:AB_476697). The secondary antibody conditions were donkey anti-rabbit (GE Healthcare, catalog number NA934, RRID:AB_772206) or sheep anti-mouse (GE Healthcare, catalog number NXA931, RRID:AB_772209) antibody (1:10000).

### Cell viability assays

Cell viability assays were carried out as described previously (38). Briefly, DOX-inducible Cas13d cells with constitutive non-targeting control or METTL1 guide 2, 4 or 5 in BE(2)-C, TN-2, Rh30, RD, SJSA-1, OZ-SBT-1476 and AUS-DIPG-017 cells were plated in media containing DMSO control or DOX [1 µg/mL for Rh30, SJSA-1, OZ-SBT-1476 and AUS-DIPG-017, and 2 µg/mL for RD, TN-2 and BE(2)-C] at 37°C in a humidified 5% CO_2_/95% air incubator. alamarBlue reagent (Thermo Fisher Scientific) was added to the media on day 4 [Rh30, SJSA-1, OZ-SBT-1476, AUS-DIPG-017, RD and BE(2)-C] or day 7 (TN-2) after DOX treatment, and each plate was read immediately on a microplate reader at 570/595 nm (Victor3, PerkinElmer, Waltham, MA) or 544/590 nm (Omega, BMG Labtech, Ortenberg, Germany) for the first reading. The plates were then incubated at 37°C in a humidified 5% CO_2_/95% air incubator for 6 hours and read again. Background from the first reading was subtracted from the second reading, and the number of viable cells was normalized to the vehicle-treated control and expressed as a percentage relative to the no-guide, DOX-treated Cas13d cells.

For the METTL1 silencing and treatment with the anti-apoptotic protein inhibitors assays, the DOX-inducible Cas13d with the constitutive non-targeting control or METTL1 guide 4 or 5 Rh30, TN-2 and BE(2)-C cells were plated in media containing DMSO control or DOX [0.5 µg/mL for Rh30, and 1 µg/mL for TN-2 and BE(2)-C; half the concentration used in the knockdown experiments] and incubated at 37°C in a humidified 5% CO_2_/95% air incubator. The next day, the cells were treated with vehicle control or a series of concentrations of ABT263 (navitoclax), S63845 or ABT199 (venetoclax) for 48 or 72 hours. alamarBlue reagent was added and plates were read twice as described above. The number of viable cells were normalized to the respective guide treated with vehicle control then expressed as a percentage of cells expressing the non-targeting control guide treated with DOX.

For the METTL1 silencing, and METTL1 and WDR4 over-expression rescue assays, each stable line was plated in triplicate wells in a 96 well plate, with eight identical plates set up. Twenty-four hours later, alamarBlue was added to one of the plates (day 0) and read on a microplate reader as described above and the rest of the plates were treated with 1 µg/mL DOX. For each subsequent day, alamarBlue was added to one of the remaining plates and read on a microplate reader. Media was changed with fresh DOX for the plates beyond 3 days of incubation time. The number of viable cells were normalized to the plate read on day 0 and expressed as a fold change of day 0.

### Clonogenic assays

The DOX-inducible Cas13d with the non-targeting control or METTL1 guide 2, 4 or 5 BE(2)-C (1000 cells/well), TN-2 (8000 cells/well), RD (1000 cells/well) or Rh30 (2000 cells/well) stable cells were plated in 6-well plates, and treated the next day with vehicle control or 2 µg/mL [BE(2)-C, TN-2 & RD] or 0.5 µg/mL (Rh30) of DOX for 10 [BE(2)-C, Rh30 & RD] or 14 (TN-2) days. During the 10- or 14-day incubation time, DOX was refreshed every 3-4 days by changing the media with fresh DOX for BE(2)-C, Rh30 and RD. Media was not changed for TN-2 due to the semi-adherent nature of the cells, but DOX was topped up every 3-4 days. The colonies were stained with 0.5% crystal violet containing 50% methanol and destained with sterile water. The colony area was quantified using Image J and normalized to vehicle control treated and expressed as a percentage relative to the no guide and DOX treated Cas13d cells to account for the effects of Cas13d alone.

### RNA-sequencing

BE(2)-C cells expressing DOX-inducible Cas13d and constitutive non-targeting control, or METTL1 guide 2, 4 or 5 were treated with vehicle control or 2 µg/mL DOX for 96 hours; DOX was refreshed every 24 hours by replacing the medium with fresh DOX-containing medium. RNA was extracted using the RNeasy Plus kit (Qiagen, Hilden, Germany), and libraries were prepared using the Illumina Stranded Total RNA Prep with Ribo-Zero Plus kit (Illumina, San Diego, CA), followed by whole-transcriptome sequencing on a NovaSeq 6000 to generate 100 million 150-bp paired-end reads (deposited in GEO, accession number GSE298373). Rh30 cells expressing DOX-inducible Cas13d and constitutive non-targeting control, or METTL1 guide 2, 4 or 5 were treated with vehicle control or 1 µg/mL DOX for 8, 24 or 48 hours; DOX was refreshed every 24 hours by replacing the medium with fresh DOX-containing medium. RNA was extracted using the RNeasy Plus kit (Qiagen), and libraries were prepared using the Illumina Stranded Total RNA Prep with Ribo-Zero Plus kit (Illumina), followed by whole-transcriptome sequencing on a NovaSeq X Plus to generate 50 million 150-bp paired-end reads (deposited in GEO, accession number GSE298686).

### RNA-sequencing data analyses

FASTQ files were pseudo-aligned to a reference index of the human transcriptome (GRCh38.p13, Ensembl release 104, GENCODE release 38, RRID:SCR_014966) using kallisto (v0.46.1) with 100 bootstraps. Integrated normalization and differential expression analysis were performed using sleuth (v0.30) with significance determined by likelihood ratio test. A significance cutoff of qval < 0.01 was used. For the Rh30 cell line, PCA analysis of the sleuth output transcript-level data identified an outlier (Rh30_g5_Ctrl 48h_repeat 3) based on the interquartile range method (i.e. exceeded the third quartile + 1.5× the interquartile range for PC1). This sample was omitted prior to re-running the kallisto-sleuth analysis pipeline. Transcripts that were found to be differentially expressed between non-targeting control (NTC) treated with DOX and NTC treated with vehicle control (Ctrl) (log_2_ fold-change <-1 or >1 and qval < 0.01) were filtered and excluded from all subsequent analyses as they are not the effects of METTL1 silencing but the effects of DOX treatment and Cas13d induction. Gene set enrichment analysis was performed against the MSigDB human hallmark gene set on significantly differentially expressed canonical transcripts ranked by log_2_ fold-change using the fgsea package (v1.28.0) in R (v4.3.1).

### METTL1 and WDR4 over-expression

METTL1 wild type (NM005371.6) open reading frame was codon optimized at the METTL1 guide 2 target sequence with 5 mismatches without altering the amino acid sequence. This was linked to T2A mApple and custom cloned into a lentiviral construct containing a neomycin resistant gene by VectorBuilder. The METTL1 mutant construct also contains the 5 mismatches with an additional point mutation at the 488^th^ base position, changing the base from A to C which results in a D163A mutation in the METTL1 protein coding sequence and deactivates the catalytic activity of METTL1. WDR4 (NM_018669.6) open reading frame linked to P2A-eGFP was also custom cloned into a lentiviral vector containing bleomycin resistant gene by VectorBuilder. Empty vector control or METTL1 wild type or mutant and WDR4 over-expression constructs were co-transfected with the pCMV-VSV-G envelope and pCMV-dR8.2 packaging plasmids into HEK293T cells using lipofectamine LTX (Invitrogen). Viral medium containing the empty vector was harvested and used to transduce the Rh30 cells expressing DOX-inducible Cas13d and non-targeting control guide. Viral medium containing the empty vector or METTL1 wild type + WDR4 or METTL1 mutant + WDR4 was harvested and used to transduce the Rh30 cells expressing DOX-inducible Cas13d and METTL1 guide 2. Cells were selected with 20 µg/mL bleomycin for 3 days followed by 1 mg/mL G418 for 6 days.

### METTL1 guide 5 RNAs packaged in lipid nanoparticles (LNPs)

For synthesizing non-targeting control (NTC) or METTL1 guide 5 crRNA packaged lipid nanoparticles, ionizable lipid (SM102), sterol lipid (β-sitosterol), helper lipid (Distearoylphosphatidylcholine or DSPC), and PEGylated lipid (1,2-Dimyristoyl-rac-glycero-3-methoxypolyethylene glycol-2000 or DMG-PEG 2000) were combined at 50:38.5:10:1.5 %mol (38). The particles were synthesized using microfluidic mixing of the organic phase containing the lipids and mRNA-containing aqueous phase (30 mM sodium acetate buffer, pH 4), using the Spark NanoAssemblr microfluidic system (Precision NanoSystems). mRNA encapsulation efficiency and concentration of the synthesized batch of LNPs was determined using the Quant-iT RiboGreen RNA assay (Invitrogen, R11490) as per manufacturer’s protocol.

### Puromycin incorporation assays

Rh30 or TN-2 with DOX-inducible Cas13d cells were plated in media containing vehicle control or DOX (1 µg/mL for Rh30 and 2 µg/mL for TN-2) and incubated at 37°C overnight in a humidified 5% CO_2_/95% air incubator. The next day, NTC or METTL1 g5 crRNA guide packaged in LNPs were added to the cells at a final concentration of 1 ng/µL and the cells were incubated at 37°C overnight in a humidified 5% CO2/95% air incubator. Media was changed the next day with fresh DOX for Rh30, but not for TN-2 due to the semi-adherent nature of the TN-2 cells and the cells were incubated 37°C overnight in a humidified 5% CO_2_/95% air incubator. Forty-eight hours after LNPs treatment, cells were pulsed with 10 µg/mL puromycin (15 minutes for Rh30 and 30 minutes for TN-2), followed by protein extraction and western blotting to assess METTL1 silencing and the effects on global translation.

### Polysome fractionation

Rh30 cells expressing DOX-inducible Cas13d together with a constitutively expressed NTC, METTL1 g2 or g5 crRNA were treated with vehicle control or 1.25 µg/mL DOX for 48 hours, reaching 70-80% confluence (10-20 × 10^6^ cells per condition). Cells were then treated with 100 µg/mL cycloheximide (CHX; C7698, Sigma) for 10 minutes at 37°C, harvested by trypsinization, pelleted (800 × g), washed in ice-cold PBS containing 100 µg/mL CHX and snap-frozen on dry ice. For polysome fractionation, pellets were lysed in buffer containing 20 mM Tris, pH 7.4, 140 mM KCl, 5 mM MgCl_2_, 1% Triton X-100, 1 mM DTT, 100 µg/ml CHX, 1 mg/mL heparin, 25 U/mL Turbo DNase I (04716728001, Roche), SUPERaseIn RNase inhibitor (AM2694, Ambion), and protease inhibitors (04693132001, Roche). Lysates were clarified at 20,000 × g for 20 minutes at 4°C. Supernatant (200 µL, 400 µg total RNA) was layered onto a 20-60% sucrose gradient in the same buffer without Triton X-100, DNase, or heparin and centrifuged at 247,600 × g in an SW 41 Ti rotor (331362, Beckman Coulter) for 3 hours and 30 minutes at 4 °C. Gradients were fractionated on a Density Gradient Fractionation System (BioComp) at 0.20 mm/sec with A_260_ detection (500 ms integration, 700 µL fractions). Profiles were all aligned to the vehicle-treated NTC sample and baselined-subtracted. Areas under the 80S and polysome regions were integrated and expressed as a ratio. For illustrative purposes, the traces were scaled to align their 80S peaks. Polysome fractionation was performed at the BioCode: RNA to Proteins Core Facility of the University of Geneva (https://www.unige.ch/medecine/r2p/R2P).

### RNA sequencing of polysome-associated RNA and total RNA

Total RNA was extracted from input and pooled polysome fractions. For input, 50 µL of whole cell lysate was processed using QIAzol Lysis Reagent (79306, Qiagen) and Direct-zol RNA Miniprep Kit (R2052, Zymo Research). Fractions containing polysomes (≥ 2 ribosomes) were pooled; RNA was extracted using QIAzol Lysis Reagent and chloroform, followed by alcohol precipitation. Libraries were prepared with the Illumina Stranded mRNA Prep, Ligation kit and sequenced on an Illumina Novaseq 6000 to a depth of approximately 50 million 100-bp single-end reads per sample (deposited in GEO, accession number GSE348070).

### Analysis of RNA sequencing of polysome-associated and total RNA

Sequencing data from total RNA and polysome fractions were quantified using the nf-core/rnaseq pipeline v3.22.2. Briefly, reads were aligned to the hg38 human genome with STAR (RRID:SCR_004463), and transcript abundance was estimated using Salmon. Salmon transcripts were filtered to retain only those belonging to protein-coding genes, and these were aggregated to gene-level counts using tximport (v1.26.1) before being used as input to DESeq2 (v1.38.3) (RRID:SCR_000154). Lowly expressed genes were removed by retaining only genes with median normalized counts > 50 in at least one experimental condition. Principal component analysis of variance-stabilized counts identified three outliers (NTC and g2 treated with vehicle control, and g2 treated with DOX, all polysomal RNA from replicate 4), which were excluded from subsequent analyses. A negative-binomial model was fitted, and contrasts were computed comparing g2, g5, or the average of g2 and g5 against NTC within either the total or polysomal fraction. Translational shift was assessed as an interaction contrast, log₂(KD polysome RNA/KD total RNA) − log₂(NTC polysome RNA/NTC total RNA). Significance was assessed by Wald test with Benjamini-Hochberg correction, applying a threshold of FDR < 0.05.

For each gene, codon usage was analyzed by calculating the fraction of each codon (codon count divided by total codon count), thereby controlling for coding sequence length. Mean and median codon fractions – including codons decoded by m^7^G-modified tRNAs as defined by Orellana et al. (19), were compared between genes with significantly reduced and significantly increased translation efficiency, and against the global mean and median. Simple linear regression through the origin was then fitted separately for codons decoded by m^7^G-modified and non-m^7^G-modified tRNAs, and slopes were compared by F-test using the car R package (v3.1.5).

## RESULTS

### *METTL1* amplification and overexpression are associated in a subset of pediatric cancers

The ZERO Childhood Cancer program undertakes whole genome sequencing (WGS) and transcriptome sequencing (TS) on all children with cancer in Australia (39). To understand the range of tumors with *METTL1* gene amplification and/or expression, we analyzed copy number and expression levels (transcripts per million, TPM) for 2099 tumor samples from ZERO (Fig. 1A and B; Supplementary Table S1). *METTL1* amplification and elevated gene expressions were observed across four main tumor types, high grade glioma (HGG), neuroblastoma (NBL), osteosarcoma (OST), and both fusion negative and positive rhabdomyosarcoma (RMS-FN, RMS-FP) (Fig. 1A). A positive correlation between *METTL1* gene copy number and METTL1 gene expression was observed, heavily influenced by the high copy number samples (Fig. 1B). CRISPR-Cas9 dependency screens in 1178 cell lines from the CRISPR 24Q4+Score Chronos dataset in the Cancer Cell Line Encyclopedia in DepMap (40) showed a selective METTL1 dependency in cell lines that include rhabdomyosarcoma and neuroblastoma (Fig. 1C).

**Fig. 1.**
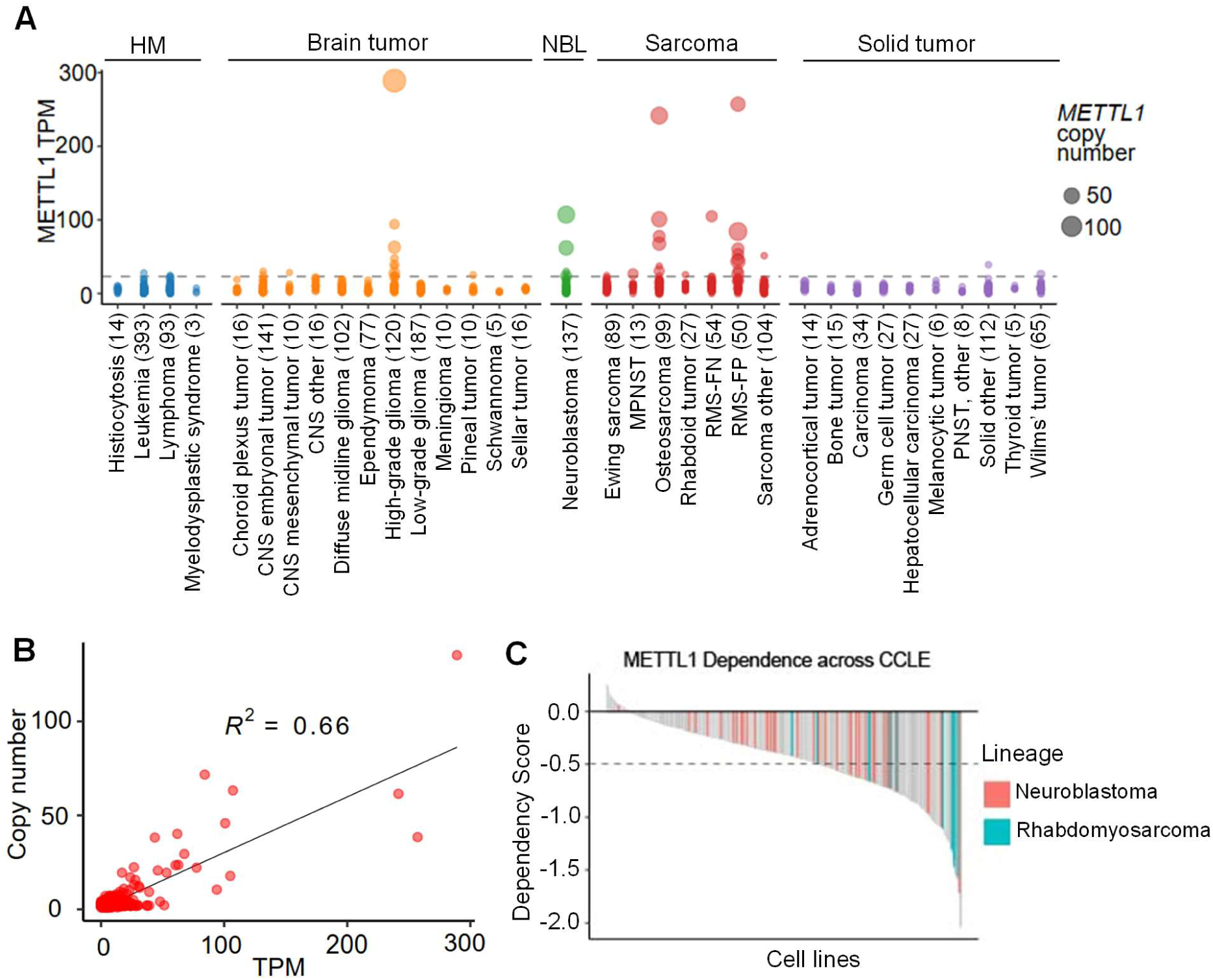
*METTL1* amplification and METTL1 overexpression are associated in pediatric cancer. (**A**) *METTL1* gene copy number and gene expression were examined in the ZERO dataset consisting of 2099 pediatric cancer patients. The horizontal dotted line marks the TPM cut-off point of quartile 3 + 3×Interquartile range. HM, hematological malignancies; CNS, central nervous system; MPNST, malignant peripheral nerve sheath tumor; PNST, peripheral nerve sheath tumor; RMS-FN, rhabdomyosarcoma fusion negative; RMS-FP, rhabdomyosarcoma fusion positive; NBL, neuroblastoma. Numbers inside parentheses indicate the number of patients. (**B**) Simple linear regression analysis showing the correlation of *METTL1* gene copy number and gene expression. (**C**) METTL1 dependency was examined in the CRISPR knockout studies in 1178 cell lines (CRISPR 24Q4+Score Chronos) from the Cancer Cell Line Encyclopedia (CCLE). Cell lines with a dependency score < -0.5 are considered dependent on METTL1 for survival.

### CRISPR-Cas13d targeting effectively reduces METTL1 expression

To determine the consequences of METTL1 RNA silencing, we selected two *METTL1* amplified [Rh30 (RMS-FP) and TN-2 (NBL)] and two *METTL1* non-amplified [RD (RMS-FN) and BE(2)-C (NBL)] cell lines (Supplementary Fig. S1) for CRISPR Cas13d-dependent METTL1 targeting. BE(2)-C, whilst not represented in DepMap, is derived from SK-N-BE(2) (Supplementary Fig. S1). We established stable doxycycline (DOX)-inducible Cas13d expression in the four selected cell lines (see Materials and Methods). Eight METTL1 CRISPR Cas13d guides (crRNA) targeting different regions of the METTL1 canonical transcript were tested, four located within exonic regions, two targeting exon-exon junctions and two targeting the exon-intron junctions (Supplementary Fig. S2A). To select the most efficient crRNAs, BE(2)-C cells expressing the DOX-inducible Cas13d were transduced individually with each of the METTL1 guides and silencing efficiency was measured by RT-PCR and western blot (Supplementary Fig. S2B and S2C). The crRNAs METTL1_g2 (g2) targeting exon 2, METTL1_g4 (g4) targeting the exon 1-2 junction and METTL1_g5 (g5) targeting exon 4 were most effective at reducing *METTL1* mRNA and protein (Supplementary Fig. S2B and S2C). These guides were selected for testing in the four selected cell lines.

Rh30, RD, BE(2)-C and TN-2 cells expressing the DOX-inducible Cas13d were transduced with a non-targeting control (NTC) crRNA or METTL1-targeting crRNAs g2, g4 or g5. RT-PCR was used to measure *METTL1* mRNA abundance. Two RT-PCR assays were designed, one employing sequence-specific primers flanking the homology regions of g2 and g4 (“amplicon 1”), and the other pair flanking the homology region of g5 (“amplicon 2”) (Fig. 2A). This was done as the proximity to the Cas13d cleavage site may alter the fidelity of these assays (41). g5 resulted in the greatest depletion of *METTL1* mRNA across all lines as detected in assays using amplicon 1 or 2 (Fig. 2B). *METTL1* mRNA silencing was accompanied by concordant decreases of METTL1 protein, demonstrated in western blots of each cell line (Fig. 2C; Supplementary Fig. S3). The METTL1 crRNAs also reduced expression of the phosphorylated form of METTL1 (AKT-dependent S27 phosphorylation (42)) (Supplementary Fig. S4).

**Fig. 2.**
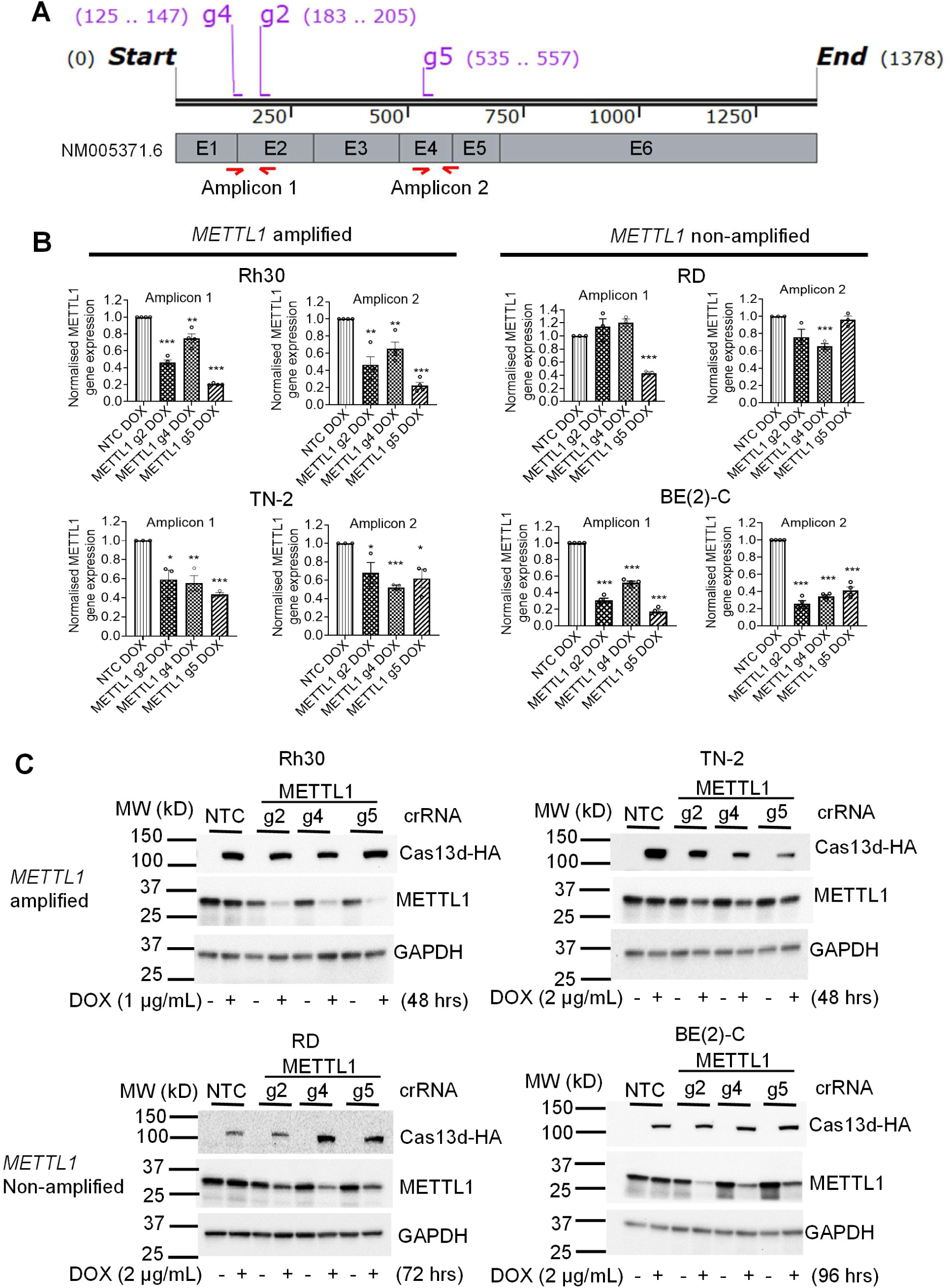
CRISPR-Cas13d targeting effectively reduces METTL1 expression. (**A**) A schematic showing the target regions of METTL1 crRNAs relative to the cDNA of METTL1 transcript NM005371.6. Numbers in brackets next to each guide crRNA indicate the position of its guide binding site. Amplicons 1 and 2 are PCR products from primer pairs 1 and 2 respectively, indicated by red arrows. The schematic is not drawn to scale. E, exon. (**B-C)** Cells expressing DOX-inducible Cas13d and a single constitutively expressed crRNA (NTC or METTL1 g2, 4 or 5) were treated with vehicle control or DOX for 48 (Rh30 and TN-2), 72 (RD) or 96 [BE(2)-C] hours, followed by RT-PCR (B) and western blot (C) analyzes. Data are presented as mean ± s.e.m. *, ** & *** indicate *P* < 0.05, 0.01 & 0.001 respectively by comparing to NTC using unpaired T-test. Replicates and densitometry analyses of the western blots are shown in Supplementary Fig. S3.

### Cas13d-dependent METTL1 silencing reduces survival and proliferation of *METTL1* amplified cell lines

We next evaluated the degree to which METTL1 silencing altered cell survival and colony formation. Knocking down METTL1 expression with either g2, g4 or g5 significantly decreased cell viability (Fig. 3A) and clonogenicity (Fig. 3B) of the *METTL1* amplified Rh30 and TN-2 cells but not non-amplified RD and BE(2)-C cells (Fig. 3A and C). The reduction in cell viability and clonogenicity was greatest in Rh30 cells, perhaps reflecting more efficient silencing. It was also evident that inducing the expression of Cas13d alone or with non-targeting crRNA (NTC) was sufficient to cause a background reduction of colony size of Rh30 and TN-2 cells (Fig. 3B). However, the substantially greater reduction in proliferation and viability in the presence of METTL1 targeting guides strongly supports the conclusion that most loss of viability and colonies was specifically due to METTL1 targeting.

**Fig. 3.**
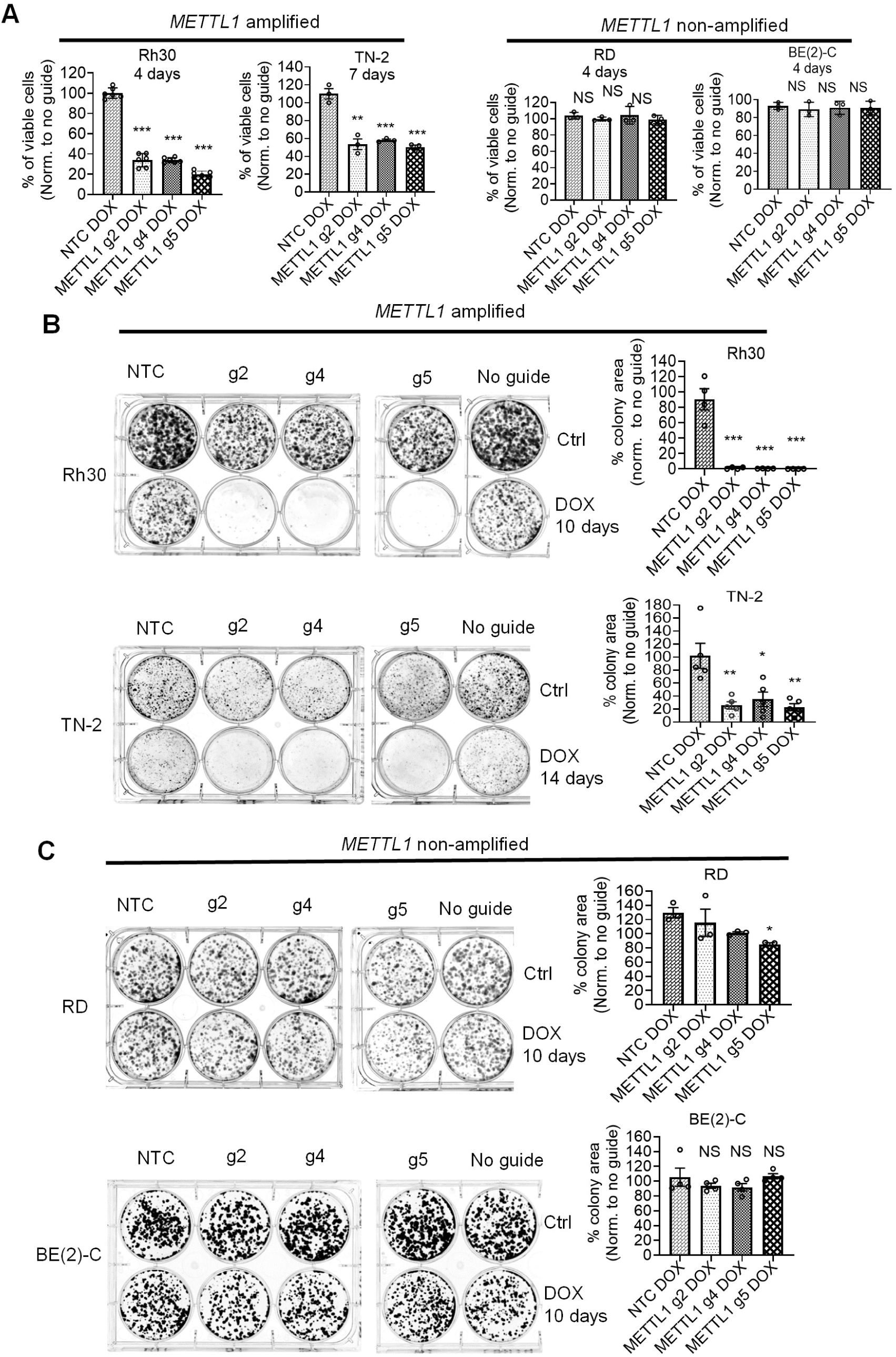
Cas13d-dependent METTL1 silencing reduces survival and proliferation of *METTL1* amplified cancer cell lines. (**A-C**) Cells expressing DOX-inducible Cas13d and a single constitutively expressed crRNA (NTC or METTL1 g2, g4 or g5) were treated with vehicle control or DOX for 4 [BE(2)-C, RD & Rh30] or 7 (TN-2) days followed by cell viability assays (A); or 10 [BE(2)-C, RD & Rh30] or 14 (TN-2) days followed by clonogenicity assays (B and C). Data are presented as mean ± s.e.m. *, ** & *** indicate *P* < 0.05, 0.01 & 0.001 respectively by comparing to NTC using unpaired T-test.

Using the most effective METTL1 crRNA, g5, we further examined the effects of METTL1 silencing in two patient-derived cell lines (OZ-SBT-1476 and AUS-DIPG-017) and the *METTL1*-amplified osteosarcoma cell line SJSA-1. OZ-SBT-1476 was established from an osteosarcoma patient (ZCCS1476) enrolled in the ZERO program (*METTL1* amplified; patient 432, Supplementary Table S1) and AUS-DIPG-017 from a patient with Diffuse Intrinsic Pontine Glioma (DIPG) (*METTL1* non-amplified) (35). g5 reduced METTL1 protein expression in all three cell lines (Supplementary Fig. S5A) and decreased cell viability in the *METTL1-*amplified osteosarcoma cell lines but not in the *METTL1* non-amplified DIPG line (Supplementary Fig. S5B).

We tested an alternative delivery method of the METTL1 guide by packaging NTC or g5 into Lipid Nanoparticles (LNPs) with a formulation shown previously to be effective in delivering crRNA (38). g5 LNPs were added to Rh30 and TN-2 cells expressing the DOX-inducible Cas13d resulting in an ∼80% reduction of METTL1 gene expression in Rh30, and 50% in TN-2 (Supplementary Fig. S6A). Similar reductions in the levels of METTL1 protein were observed (Supplementary Fig. S6B). This delivery system also reduced cell viability, but less than lentiviral delivery methods, likely as a result of the shorter half-life of LNP-delivered crRNAs (Supplementary Fig. S6C). Nevertheless, this alternative delivery system further demonstrates the METTL1 dependency of *METTL1*-amplified cell lines.

### Rescuing METTL1 silencing requires METTL1 catalytic activity

The oncogenic activity of overexpressed METTL1 in some tumor types is not solely dependent on METTL1 catalytic activity (25). Structural and functional analyses show that 3 amino acid residues in the catalytic domain of METTL1 (E240, D199 and D163) are critical for methylation activity and mutations of any of these residues (to alanine) abolish the catalytic activity (42). We exploited this to investigate the requirement of the methyltransferase activity of METTL1 in METTL1 dependency, and designed and cloned codon optimized METTL1 and a catalytically inactive mutant (D163A) open reading frames which harbored 5 base-pair mismatches to METTL1 g2 crRNA, but still permitted the translation of METTL1 (Fig. 4A). These 5 base-pair mismatches are beyond the maximum mismatch tolerance of RfxCas13d (34), and are expected to abolish the silencing activity against the ectopically expressed codon-optimized METLL1, but not the endogenous METTL1. This approach also required the ectopic expression of WDR4, the interacting partner required for METTL1 activity (Fig. 4B and C). Using this system, we compared the capacity of catalytically active and inactive forms of METTL1 to rescue cells from cell death induced by g2-dependent silencing of METTL1. Western blot analysis showed that g2 expressing Rh30 cells transduced with METTL1 wild type and WDR4 or METTL1 mutant and WDR4 overexpression constructs have exogenous WDR4 and METTL1 expression and diminished endogenous METTL1 (Fig. 4B). Cell viability assays showed that silencing of endogenous METTL1 by g2 induced cell death that could only be partially rescued by the co-expression of catalytically active METTL1 and WDR4 (Fig. 4C). These data indicate that the catalytic activity of METTL1 is required for the viability of Rh30 cells, and consequently that the catalytic activity of METTL1 is important to METTL1 oncogenic activity in this model.

**Fig. 4.**
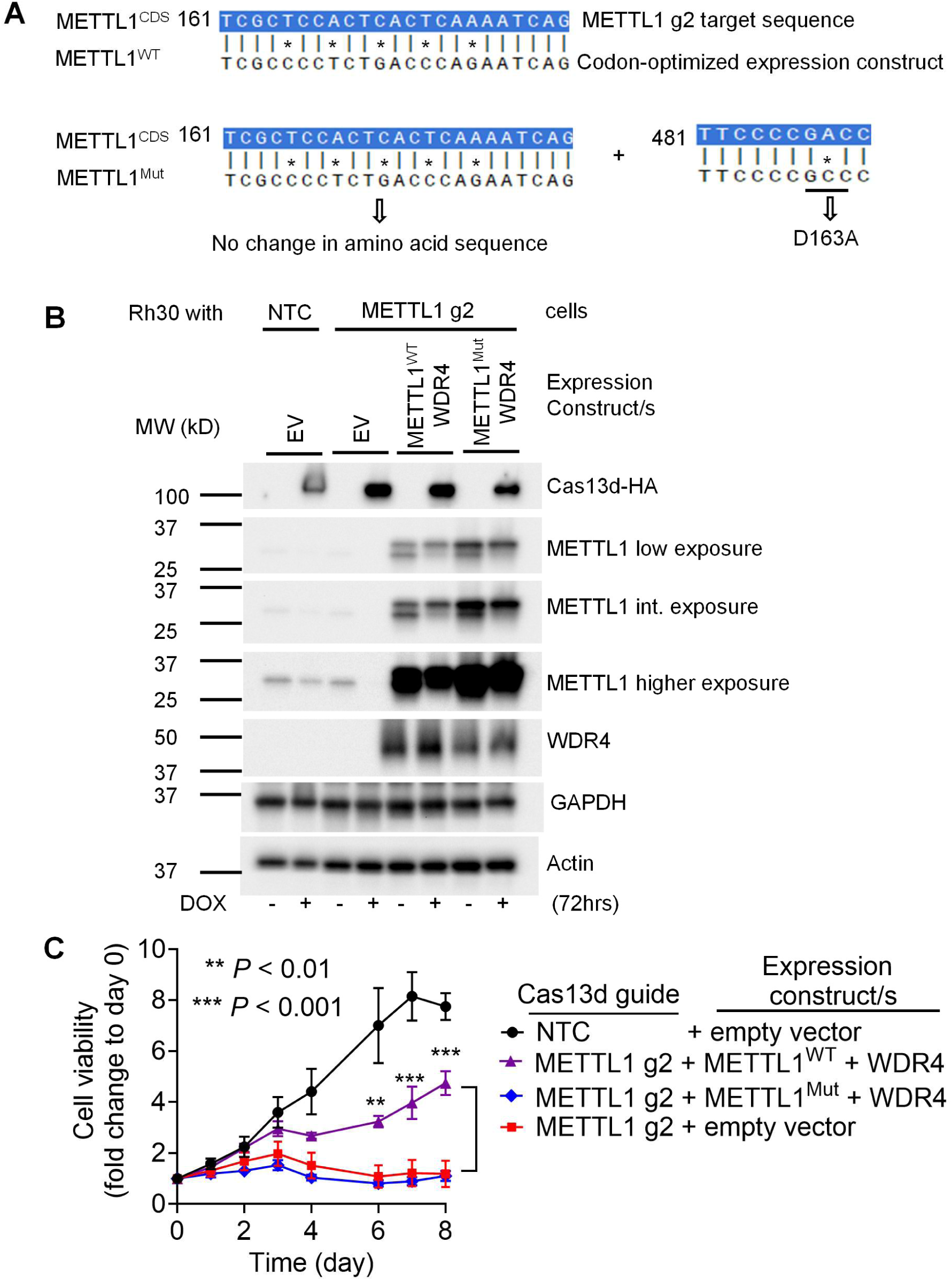
Rescuing METTL1 silencing requires METTL1 catalytic activity. (**A**) A schematic of codon optimization at the METTL1 g2 crRNA target region in both the wild type (WT) and catalytic mutant (Mut) METTL1 open reading frame constructs, and an additional single point mutation in the catalytic mutant construct. Numbers indicate the position of the NM_005371.6 METTL1 transcript coding sequence (CDS). (**B-C**) Rh30 cells expressing DOX-inducible Cas13d cells and constitutive NTC and empty vector construct or METTL1 guide 2 crRNA and empty vector construct or METTL1 guide 2 crRNA and METTL1^WT^ plus WDR4 ORF constructs or METTL1 guide 2 crRNA and METTL1^Mut^ plus WDR4 ORF constructs were treated with vehicle control or DOX (1 µg/mL) for the indicated time followed by western blot analysis (B) or cell viability assays (C). Data are presented as mean ± s.e.m. ** and *** indicate *P* < 0.01 and 0.001 respectively by comparing to METTL1 g2 crRNA plus empty vector using unpaired T-test.

### Transcriptional consequences of *METTL1* mRNA silencing are specific

To ascertain the specificity of Cas13d-dependent METTL1 silencing and the broader impacts on the transcriptome, we harvested RNA from BE(2)-C and Rh30 cells harboring Cas13d and either the non-targeting or targeting guides, with and without DOX treatment. Total RNA was collected at three time points over a 48-hour period in the METTL1-dependent Rh30 cells, a window that precedes the onset of proliferative arrest and hence captures transcriptional changes directly from METTL1 loss rather than secondary stress response. In the METTL1 independent BE(2)-C cells, which show no fitness defect upon METTL1 silencing, RNA was collected at 96 hours to allow any subtler transcriptional changes to accumulate (Fig. 5A; Supplementary Fig. S7A). There were no significant batch effects across the 4 replicate experiments (Supplementary Fig. S7B; Supplementary Fig. S8A and S8B). One outlier sample (Rh30 g5 Ctrl, 48h, replicate 3) (Supplementary Fig. S8A) was excluded on PCA analysis (Supplementary Fig. S8B). One further feature of note was the apparent accumulation of a non-coding METTL1 transcript (ENST00000549773.1) in the presence of Cas13d and g2, g4 or g5. This transcript retains intron 2 of METTL1 (Supplementary Fig. S7C and S7E; Supplementary Fig. S8C and S8D) and likely represents an accumulation of transcripts retaining intronic and non-coding sequences which arise during the mRNA degradation process (43).

**Fig. 5.**
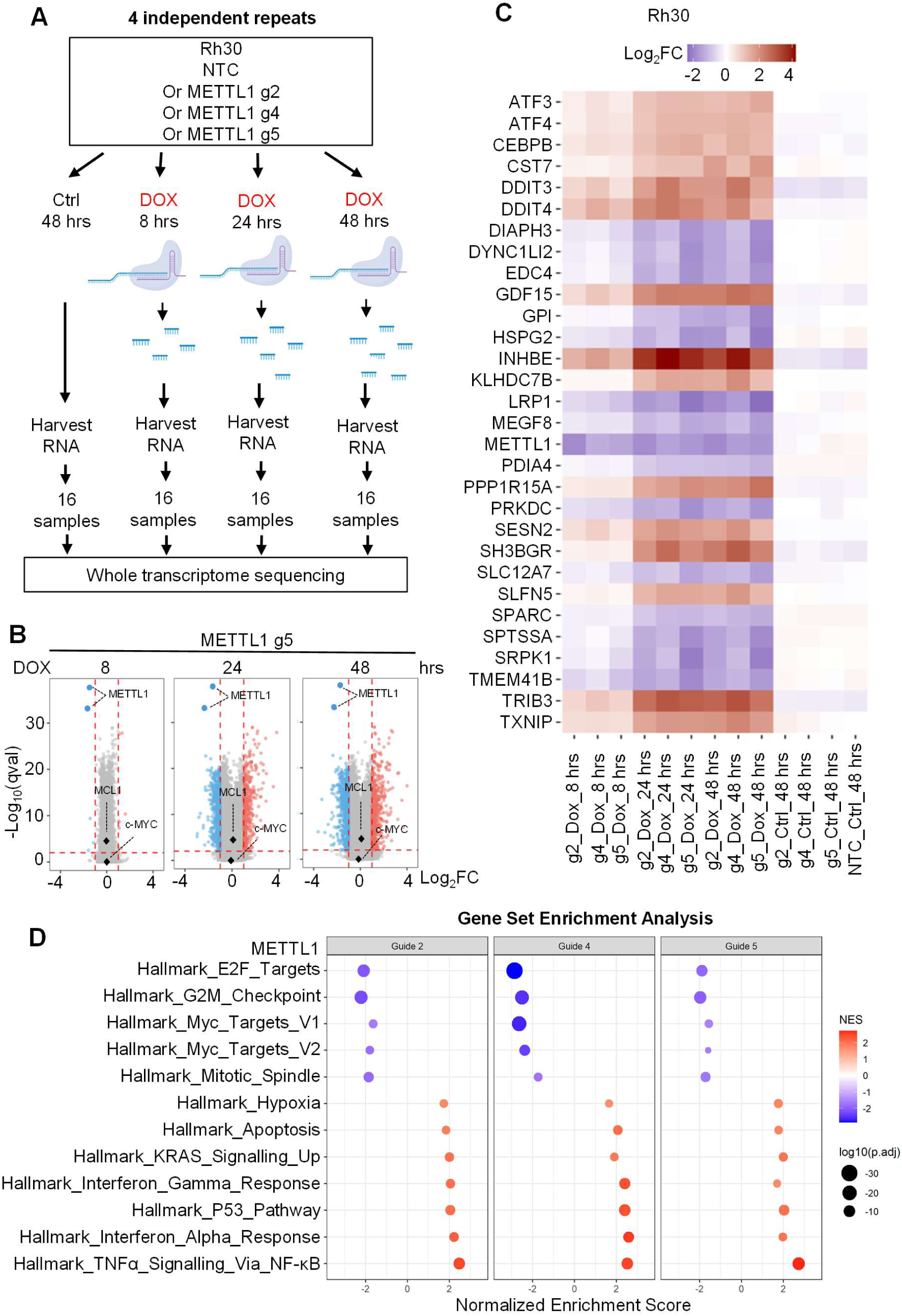
Transcriptional consequences of *METTL1* mRNA silencing are specific. (**A**) A schematic of experimental design for whole transcriptome sequencing after METTL1 silencing in Rh30 cells. (**B**) Transcriptome wide differential transcript expression (DTE) analysis showing the common transcripts up- or down-regulated in four independent experimental repeats as volcano plots, only g5 is shown (g5 is shown again with g2 and g4 in Fig. S8C). The vertical dotted lines mark the cut-off points using log_2_ fold-change <-1 or >1 and the horizontal dotted line marks the cut-off point using qval <0.01. Blue dots on the left of the x-axis and orange dots on the right of the x-axis represent the transcripts that are significantly down- or up-regulated respectively after METTL1 silencing. (**C**) The common top 30 most significantly differentially expressed canonical transcripts in the four independent experimental repeats and three time points are presented as a heatmap. (**D**) Gene Set Enrichment Analysis of the pathways that were significantly modulated in the 48-hour time point. NES, normalized enrichment score.

Differential transcript expression (DTE) analysis was used to identify the transcriptional consequences of *METTL1* mRNA silencing, with a log_2_ fold-change >1 or <-1 and qval < 0.01 in transcripts in each of the 4 experiments defined as a significant change in transcript abundance. Most striking was that METTL1 transcripts were the most significantly downregulated in each cell line in the presence of a METTL1 targeting guide, and the only significantly differentially expressed gene in the non-METTL1 dependent BE(2)-C cells 96 hours after METTL1 silencing (Supplementary Fig. S7C and S7D; Supplementary Table S2). In contrast, Rh30 cells showed a progressive accumulation of significant transcriptional changes over the 48-hour period (Fig. 5B; Supplementary Fig. S8C; Supplementary Table S3). The top 30 most differentially expressed canonical transcripts are shown in the heatmap in Fig. 5C. Gene Set Enrichment Analysis (GSEA) showed an upregulation of apoptosis and TP53 pathways and a downregulation of MYC targets, E2F targets, G2M checkpoint pathways and mitotic spindle pathways, all consistent with cells undergoing cell cycle arrest and cell death (Fig. 5D; Supplementary Table S4). The significance of the apparent upregulation of KRAS signalling pathway is unknown. The mechanistic implications of the upregulation of inflammatory pathway genes (interferon (IFN) alpha and gamma responses, tumor necrosis factor alpha (TNFα) signalling) in this tumor cell line are also not clear.

### Downregulation of METTL1 is associated with reduced MCL1 and MYC protein expression

Since *METTL1* mRNA silencing was associated with increased cell death and transcriptional activation of apoptosis pathways, we used western blotting to determine the impact of reducing METTL1 expression on anti-apoptotic BCL2 family members (Fig. 6; Supplementary Fig. S9 - Fig. S12). Across all experiments, MCL1 protein expression was decreased in the presence of METTL1 targeting guides and Cas13d (Fig. 6A and B; Supplementary Fig. S9 - Fig. S12) in the Rh30 cells, but less consistent changes in BCL2 and BCL-xL. Consistent with the induction of apoptosis there was more cleaved caspase-3 and PARP following the induction of Cas13d in the presence of the METTL1 targeting guides (Fig. 6A and B; Supplementary Fig. S9 - Fig. S12). Also following on from the RNA-seq analysis, c-MYC and N-MYC protein expression was determined.

**Fig. 6.**
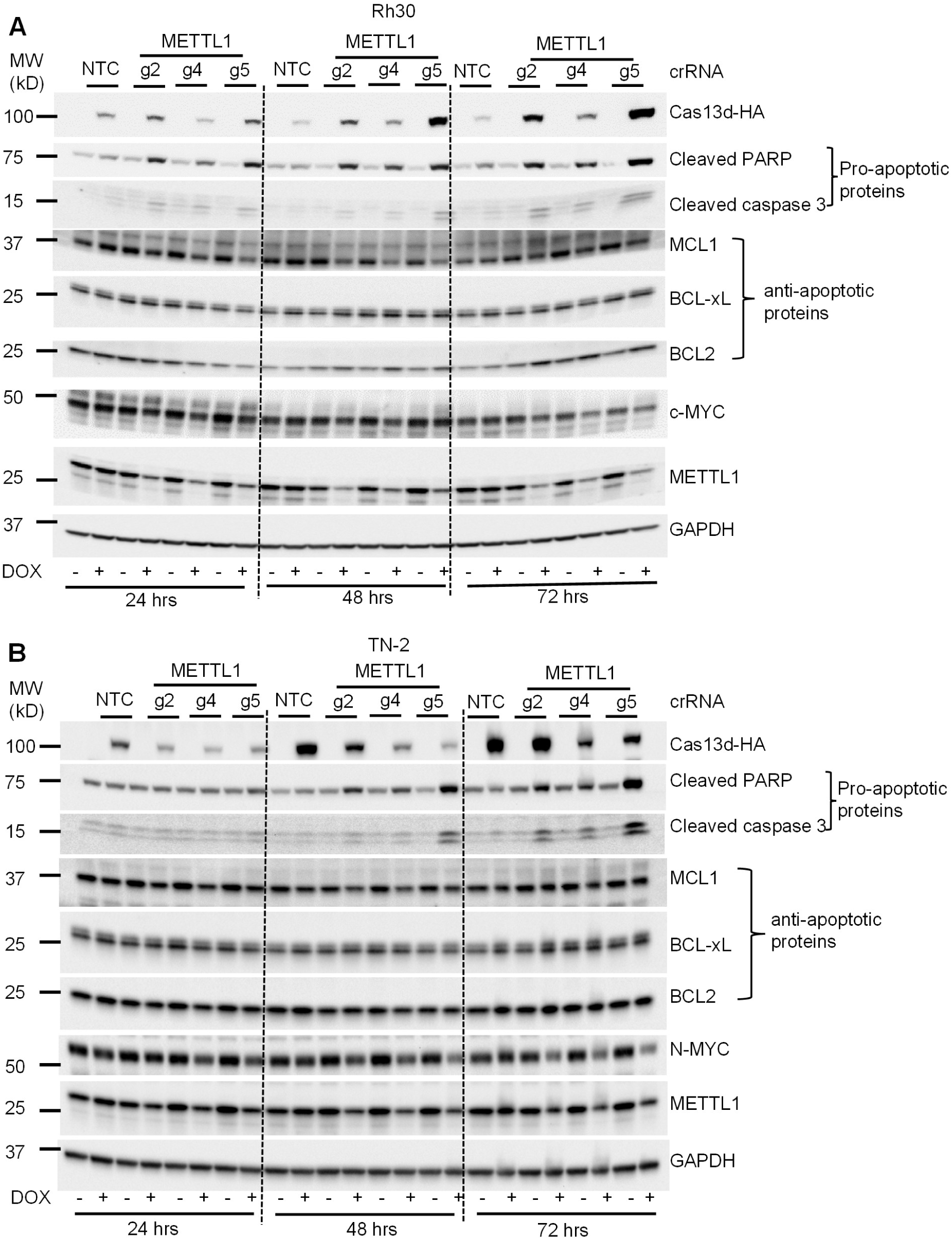
Downregulation of METTL1 is associated with reduced MCL1 and MYC protein expression. (**A-B**) Cells expressing DOX-inducible Cas13d and a single constitutively expressed crRNA (NTC or METTL1 g2, g4 or g5) were treated with vehicle control or DOX for 24, 48 and 72 hours, followed by western blot analysis of the indicated protein expression. Replicates and densitometry analyses of the western blots are presented in Supplementary Fig. S9 – Fig. S12.

After 72 hours of METTL1 silencing, c-MYC (Rh30) and N-MYC (TN-2) protein expression decreased (Fig. 6A and B; Supplementary Fig. S9 - Fig. S12). Notably, we did not observe significant changes in MCL1 and c-MYC mRNA abundance (Fig. 5B; Supplementary Fig. S8C; Supplementary Table S3). This suggests that reduced METTL1-dependent m7G methylation of tRNAs and resultant tRNA instability causes post translational reduction of MCL1 and MYC protein expression (19).

### METTL1 inhibition alters protein translation

Given the importance of METTL1 in tRNA methylation and protein translation, we examined the translational consequences of METTL1 depletion in Rh30 and TN-2 cells. We first used a puromycin incorporation assay to examine global translation. Puromycin is incorporated into newly translated protein, terminating peptide extension which can be detected using a specific puromycin antibody (44). METTL1 silencing resulted in significantly diminished puromycin incorporation when compared to cells expressing a NTC crRNA (Fig. 7A; Supplementary Fig. S13A). We performed polysome fractionation as an orthogonal method to study the effects of METTL1 on global translation in Rh30 cells expressing DOX-inducible Cas13d and NTC or METTL1-targeting guides. Similarly, METTL1 inhibition caused a significant reduction in polysome-to-monosome ratio with both g2 and g5 but not with DOX-treated NTC versus vehicle-treated NTC. (Fig. 7B and C; Supplementary Fig. S13B). Together, these data show that METTL1 silencing is associated with significant changes in translation.

**Fig. 7.**
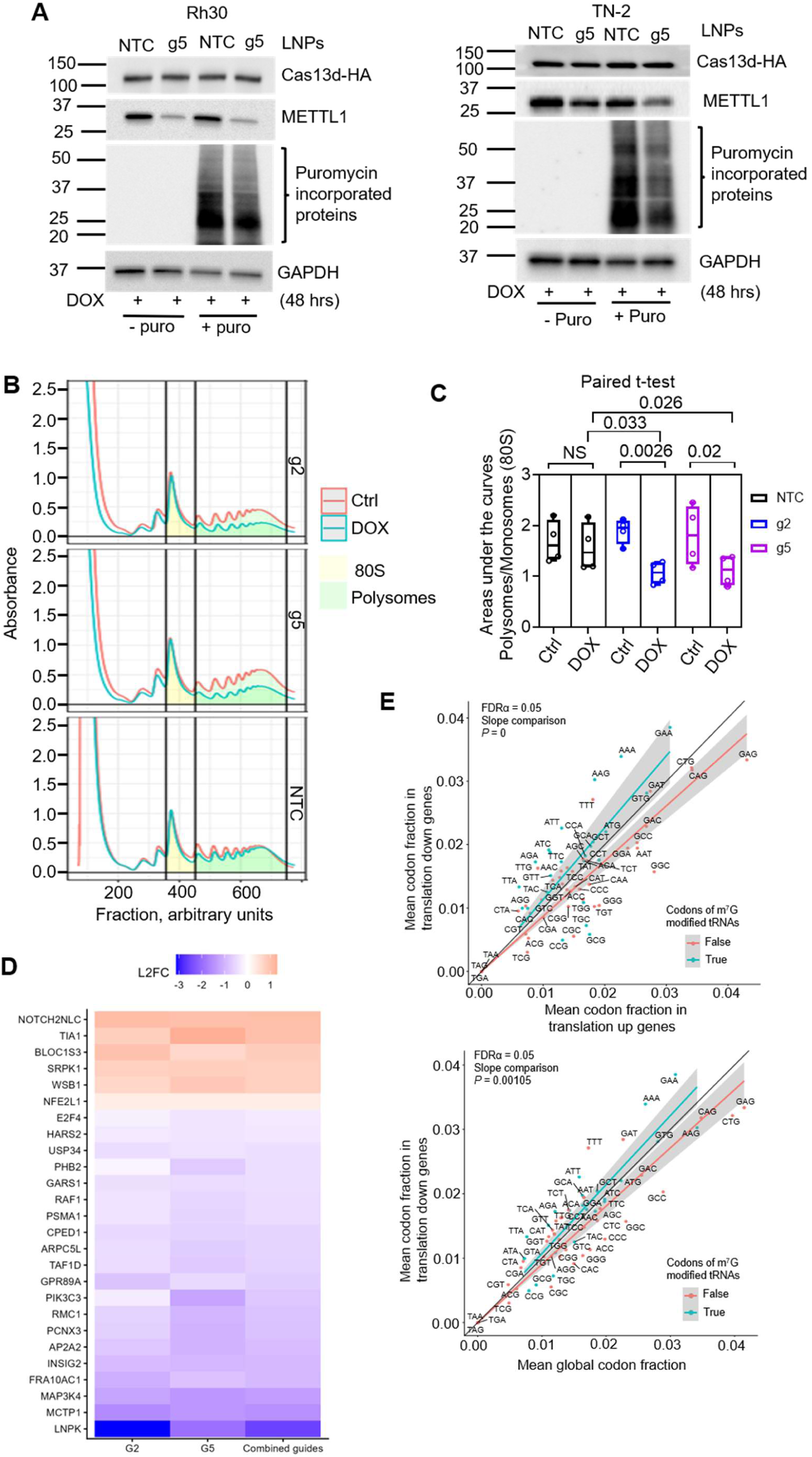
METTL1 inhibition alters protein translation. (**A**) Rh30 and TN-2 cells expressing DOX-inducible Cas13d were treated with DOX for 12 hours, followed by the addition of NTC or METTL1 g5 crRNA packaged in LNPs. Forty-eight hours post LNPs treatment, the cells were pulsed with puromycin followed by protein extraction and western blot analysis of the indicated protein expression. Replicates and densitometry analyses of the western blots are presented in Fig. S13A. (**B**-**C**) Rh30 cells expressing DOX-inducible Cas13d and constitutive NTC, or METTL1 g2 or 5 crRNA were treated with vehicle control or DOX for 48 hours, followed by treatment with cycloheximide and polysome fractionation (B). The polysome profiles (replicate 4) are presented again together with other replicates (replicate 1, 2 and 3) in Supplementary Fig. S13B. Areas under the polysomes and monosomes (80S) curves were calculated and expressed as a ratio (C). (D) The genes that are significantly differentially translated in at least three independent experimental repeats after METTL1 silencing are presented as a heatmap. (E) Mean codon usage was compared after METTL1 silencing. The upper panel compares 20 translationally downregulated genes with 6 translationally upregulated genes; the lower panel compares the same 20 downregulated genes with the global codon fraction. Each point represents a single codon, and in each panel the diagonal black line indicates equal codon usage between the two groups being compared. Blue and red lines show linear regressions for codons decoded by m^7^G-modified and non-m^7^G-modified tRNAs, respectively. Grey shaded areas correspond to the 95% confidence interval for the predicted regression.

To define the translational consequences and identify any specific transcripts selectively regulated at the translational level, we performed RNA sequencing of both total RNA and pooled heavy polysomal RNA following polysome fractionation. This approach allowed us to differentiate changes in transcript abundance from changes in translational efficiency, providing a systematic way of detecting METTL1-dependent translational programmes. Principal component analysis identified three outlier samples (NTC and g2 treated with vehicle control, and g2 treated with DOX, all polysomal RNA from replicate 4; Supplementary Fig. S14A), which were excluded from further analysis. Clustering after their removal is shown in Supplementary Fig. S14B. For the remaining samples treated with DOX, we calculated log_2_ fold changes in gene expression upon METTL1 silencing, comparing g2, g5 or the average of the guides against NTC, in both total RNA (Supplementary Fig. S14C, upper panel; Supplementary Table S5) and polysomal RNA (Supplementary Fig. S14C, middle panel; Supplementary Table S6).

Relative translation efficiency was determined using DESeq2 and expressed as the log₂ fold change in polysome association, calculated as the polysome-to-total RNA ratio for g2, g5, or the average of the two guides, divided by the corresponding ratio for NTC (Supplementary Fig. S14C, lower panel; Supplementary Table S7). Using the averaged guide comparison, we identified 20 transcripts with significantly reduced and 6 with significantly increased translation efficiency following METTL1 downregulation (Fig. 7D). Some of the translationally reduced genes have roles in oncogenesis such as RAF1 (45), which may contribute to the phenotypic effects of METTL1 silencing. However, the total decrease in polysome mRNA loading points to a more general decrease in translation accounting for METTL1-dependent cell death.

To determine whether transcripts with significantly altered translational efficiency after METTL1 silencing are enriched in codons decoded by m^7^G-methylated tRNAs, we performed codon usage analysis, comparing the mean or median codon fraction in genes with significantly reduced translation efficiency either with that in genes with significantly increased translation efficiency or with the mean or median global codon fraction (Fig. 7E; Supplementary Fig. S14D). We found that genes with decreased translation efficiency were enriched in codons decoded by m^7^G-methylated tRNAs compared with genes with increased translation efficiency, most notably GAA, AAG and AAA (Fig. 7E; Supplementary Fig. S14D). These codons are decoded by tRNA-Glu(TTC), tRNA-Lys(CTT) and tRNA-Lys(TTT), respectively, all of which carry m^7^G. This is consistent with the effects on cell growth and viability after METTL1 silencing resulting from a decrease in m^7^G methylation of a subset of tRNAs and the consequent reduction in translation.

### METTL1 inhibition synergizes with a BCL2-family protein inhibitor

We next turned to consider the consequences of diminished MCL1 protein expression after METTL1 silencing, reasoning that the reduction of MCL1 protein induced by METTL1 silencing may leave cells dependent on anti-apoptotic BCL2 family members, BCL2 and/or BCL-xL. We tested this hypothesis by silencing METTL1 expression in Rh30 and TN-2 cells, whilst simultaneously exposing cells to serial dilutions of ABT-263 (BCL-xL, BCL2 and BCL-w inhibitor), S63845 (MCL1 inhibitor) and ABT-199 (BCL2 inhibitor) (Fig. 8; Supplementary Fig. S15). BE(2)-C cells were used as a non-METTL1 dependent control. The greatest synergy was observed in both METTL1-dependent cell lines treated with the ABT-263 agent (Fig. 8A; Supplementary Fig. S15A). Weaker synergy between METTL1 silencing and the MCL1 inhibitor S63845 was observed (Fig. 8B; Supplementary Fig. S15B) and ABT-199 at high concentrations only in TN-2 cells (Fig. 8C; Supplementary Fig. S15C). Together, these data suggest that METTL1 silencing leaves METTL1-dependent cells more susceptible to agents targeting the anti-apoptotic BCL2 family, of which BCL-xL was the most important.

**Fig. 8.**
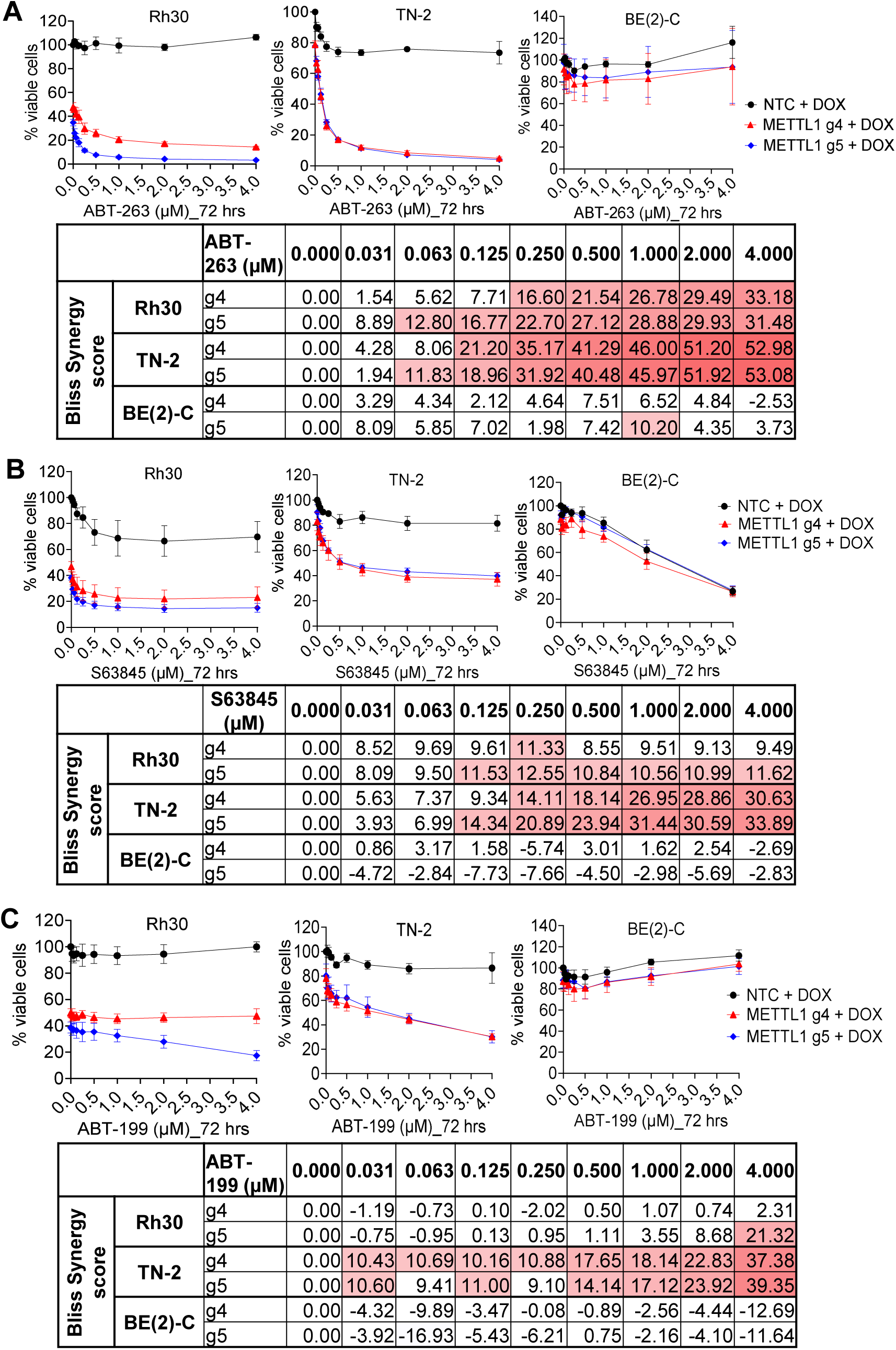
METTL1 inhibition synergizes with a BCL2 family protein inhibitor. (**A-C**) Cells expressing DOX-inducible Cas13d and a single constitutively expressed crRNA (NTC or METTL1 g4 or g5) were treated with vehicle control or DOX [0.5 µg/mL for Rh30, 1 µg/mL for BE(2)-C and TN-2; half the concentration used in knockdown cell viability assays] and ABT-263 (A), S63845 (B) or ABT-199 (C) at the indicated concentrations for 72 hours, followed by cell viability assays. Bliss synergy scores > 10 and 15 indicate synergy and strong synergy respectively.

## DISCUSSION

There is considerable interest in RNA modifying enzymes as potential dependencies and druggable targets in various cancer subtypes. In this work we focused on the role of METTL1 in extracranial pediatric solid cancers, prompted by evidence from CRISPR screens that METTL1 is a dependency gene in some rhabdomyosarcoma and neuroblastoma cell lines (40). We found that knocking down METTL1 using Cas13d-dependent mRNA degradation diminished cell viability and clonogenicity in selected rhabdomyosarcoma, osteosarcoma and neuroblastoma cell lines. This is consistent with early results reporting the effects of a first-in-class small molecule METTL1 inhibitor (29). In our selected lines, susceptibility to METTL1 silencing correlated with *METTL1* gene amplification, but this is unlikely to be the only biomarker of response because knockout screens (in sources such as DepMap) shows that some cancer cell lines with normal *METTL1* gene copy number remain dependent on METTL1 expression.

In liposarcoma, METTL1 oncogenic activity is not solely dependent on m^7^G methyltransferase activity (25). Instead, catalytically inactive METTL1 co-operates with active AKT (Protein Kinase B) to maintain the pool of aminoacylated tRNAs required for the increased translation in cancer cells. Whether the catalytic activity of METTL1 is or is not critical for the oncogenic functions of METTL1 has significant implications for the potential efficacy and application of small molecule METTL1 catalytic inhibitors. Our data suggests that in rhabdomyosarcoma cells, METTL1 catalytic activity is critical to the dependency of these cells on METTL1, because only re-expression of catalytically active METTL1 rescued cell viability after silencing of endogenous METTL1, albeit that viability was not restored to that of control cells. If catalytic inhibitors of METTL1 are to have a clinical application, it will be important to establish in which tumors the catalytic activity of METTL1 is required for METTL1 dependency. The data we present here validate the potential of inhibitors of METTL1 catalytic activity and defines potential biomarkers of response in pediatric tumors with elevated *METTL1* copy number and expression. In addition, our data provide new insights into potential therapeutic synergies with METTL1 inhibition, specifically BH3-mimetic drugs targeting anti-apoptotic BCL2-family members. The logic of this approach is supported by the loss of MCL1 expression, at least in some cell lines, which most likely results from diminished translation of short half-life proteins. METTL1 inhibition may, in this context, be thought of as priming cells for apoptosis (46). The most effective synergy was observed with the BCL2L1 inhibitor (ABT-263 or navitoclax). However, this may vary substantially across tumor types, depending on the expression and regulation of BCL2 family proteins.

There are limitations to our study that deserve further consideration. Whilst we have shown that *METTL1* mRNA degradation impacts on translation in genes with some bias for m^7^G tRNA codons, we do not know if the major impacts of METTL1 silencing are mediated through a small set of specifically METTL1-regulated genes or are a result of a generalized loss of translation of many genes. We can only speculate that the later mechanism is more likely given the substantial reduction in mRNA loading on polysomes. Differentiating between these possibilities may be informative of other rational drug combinations and potential resistance mechanisms. A second caution relates to the specificity and adaptability of the Cas13d system we deployed to silence METTL1 expression. These systems may be prone to a phenomenon called collateral cleavage. This refers to the capacity of RfxCas13d, once activated by a targeting crRNA, to indiscriminately degrade other RNAs and so contribute to any observed phenotype (47). The observed base line toxicity with Cas13d expression in the presence of both NTC and targeting crRNAs, does show that this system may cause cell loss and slow proliferation independent of specific mRNA targeting. If this were due to collateral cleavage, this would be more prominant in cells with a targeting crRNA, which was not what we observed. The most striking indication that collateral cleavage was not a major confounder in our data was the remarkable specificity of METTL1 targeting, particularly in non-METTL1 dependent cell lines. Some more recent evidence suggests collateral activity is significantly alleviated by lentiviral RfxCas13d expression as opposed to transient transfection (48). Finally, although we present data that supports the contention that the catalytic activity of METLL1 is required by METTL1-dependent cancer cell lines, we cannot and should not dismiss the possibility that the non-catalytic functions also contribute to METTL1 oncogenic mechanisms in the cell lines we studied, since the rescue experiment did not fully restore viability.

In conclusion, we have identified and validated METTL1 as a dependency in *METTL1*-amplified pediatric cancer cell lines. Although we use a genetic methodology to inhibit METTL1 activity, our approach has been remarkably specific. Our data support a model in which the inhibition of METTL1 catalytic activity may effectively induce cancer cell death, and may co-operate with agents that target the intrinsic cell death pathways. Even though we do not yet know whether METTL1-dependency is explained by diminished translation of a set specific genes or all genes, METTL1 is emerging as a legitimate anti-cancer drug target.

### Statistical analysis

Except for optimization assays and a western blot assay with multiple time points, all experiments described were performed with a minimum of three independent repeats. Statistical analyses were performed using the built-in functions in GraphPad Prism, with a two-tailed unpaired Welch’s *t*-test used to compare two groups unless otherwise specified. Error bars denote the mean ± standard error, and *P* < 0.05 was considered statistically significant.

## List of Supplementary Materials

Fig. S1. *METTL1* is amplified and overexpressed in a subset of pediatric cancer cell lines.

Fig. S2. CRISPR-Cas13d targeting effectively reduces METTL1 expression.

Fig. S3. Additional western blots and densitometry analyses show that CRISPR-Cas13d targeting effectively reduces METTL1 protein expression.

Fig. S4. CRISPR-Cas13d targeting effectively reduces the expression of phosphorylated METTL1 protein.

Fig. S5. Cas13d-dependent METTL1 silencing reduces survival of *METTL1* amplified cancer cell lines.

Fig. S6. METTL1 g5 crRNA packaged in LNPs effectively reduces METTL1 expression and cell viability.

Fig. S7. Transcriptional consequences of *METTL1* mRNA silencing are specific in BE(2)-C.

Fig. S8. Transcriptional consequences of *METTL1* mRNA silencing are specific in Rh30.

Fig. S9. Additional western blot analyses show downregulation of METTL1 is associated with reduced MCL1 and c-MYC protein expression in Rh30.

Fig. S10. Densitometry analyses of western blots show downregulation of METTL1 is associated with reduced MCL1 and c-MYC protein expression in Rh30.

Fig. S11. Additional western blot analyses show downregulation of METTL1 is associated with reduced N-MYC protein expression in TN-2.

Fig. S12. Densitometry analyses of western blots show downregulation of METTL1 is associated with reduced N-MYC protein expression in TN-2.

Fig. S13. Downregulation of METTL1 is associated with reduced global protein translation.

Fig. S14. Translational consequences of *METTL1* mRNA silencing in Rh30.

Fig. S15. METTL1 inhibition synergizes with a BCL2 family protein inhibitor.

## Other Supplementary Materials

Supplementary Table S1. *METTL1* gene expression (tpm) and copy number (cn) from tumors of 2099 pediatric cancer patients from the ZERO Childhood Cancer Program.

Supplementary Table S2. Differential gene expression analysis in BE(2)-C cells between non-targeting control and METTL1 silencing.

Supplementary Table S3. Differential gene expression analysis in Rh30 cells between non-targeting control and METTL1 silencing.

Supplementary Table S4. Gene Set Enrichment Analysis (GSEA) in Rh30 cells between non-targeting control and METTL1 silencing.

Supplementary Table S5. Differential gene expression analysis of input total RNA from Rh30 cells, collected prior to polysome fractionation.

Supplementary Table S6. Differential gene expression analysis of RNA from pooled heavy polysome fractions (≥2 ribosomes) of Rh30 cells.

Supplementary Table S7. Relative translation efficiency in Rh30 cells following METTL1 silencing.

## Acknowledgments

We sincerely thank patients and parents who participate in the ZERO Childhood Cancer study. We thank the Australian BioCommons, Velsera/SevenBridges and Children’s Hospital of Philadelphia Center for Data-Driven Discovery in Biomedicine for supporting the development of the CAVATICA genome analysis platform. The authors acknowledge the provision of computing and data resources provided by the Australian BioCommons Leadership Share (ABLeS) program. This program is co-funded by Bioplatforms Australia (enabled by NCRIS), the National Computational Infrastructure and Pawsey Supercomputing Research Center. The authors acknowledge Luminesce Alliance - Innovation for Children’s Health for its contribution and support. Luminesce Alliance is a not-for-profit cooperative joint venture between the Sydney Children’s Hospitals Network, the Children’s Medical Research Institute and the Children’s Cancer Institute. It has been established with the support of the NSW Government to coordinate and integrate pediatric research. Luminesce Alliance is also affiliated with the University of Sydney and the University of New South Wales Sydney.

## Funding

Medical Research Future Fund (MRFF EPCD000015)

Luminesce Alliance “ENP2 Translational Tumor Biology” (RG231235)

Cancer Institute New South Wales (2021/CDF1139)

National Health and Medical Research Council (Ideas Grant APP2028235)

Audi Foundation

## Author contributions

Conceptualization: PGE, PYL, IS

Methodology: PGE, PYL

Investigation: PGE, PYL, AdW, SH, AS, PR, MF, OP, FB, ZB, EF

Visualization: PGE, PYL, AdW, SH

Funding acquisition: PGE, AdW

Project administration: PGE

Supervision: PGE, PYL

Writing – original draft: PGE, PYL

Writing – review & editing: PGE, PYL, SH, AdW, AS, PR, WH, CM, MW, GMA, IS, MF, OP, FB, ZB, EF

## Data and materials availability

All data are available in the main text or the supplementary materials. The RNA sequencing raw data of this article is available at the NCBI GEO database website, with accession number GSE298373 for BE(2)-C, GSE298686 for Rh30, and GSE348070 for RNA sequencing of polysome-associated RNA and total RNA in Rh30.

