## Supplementary Figures for "METTL1 gene dosage and catalytic activity define a targetable vulnerability in pediatric cancers"

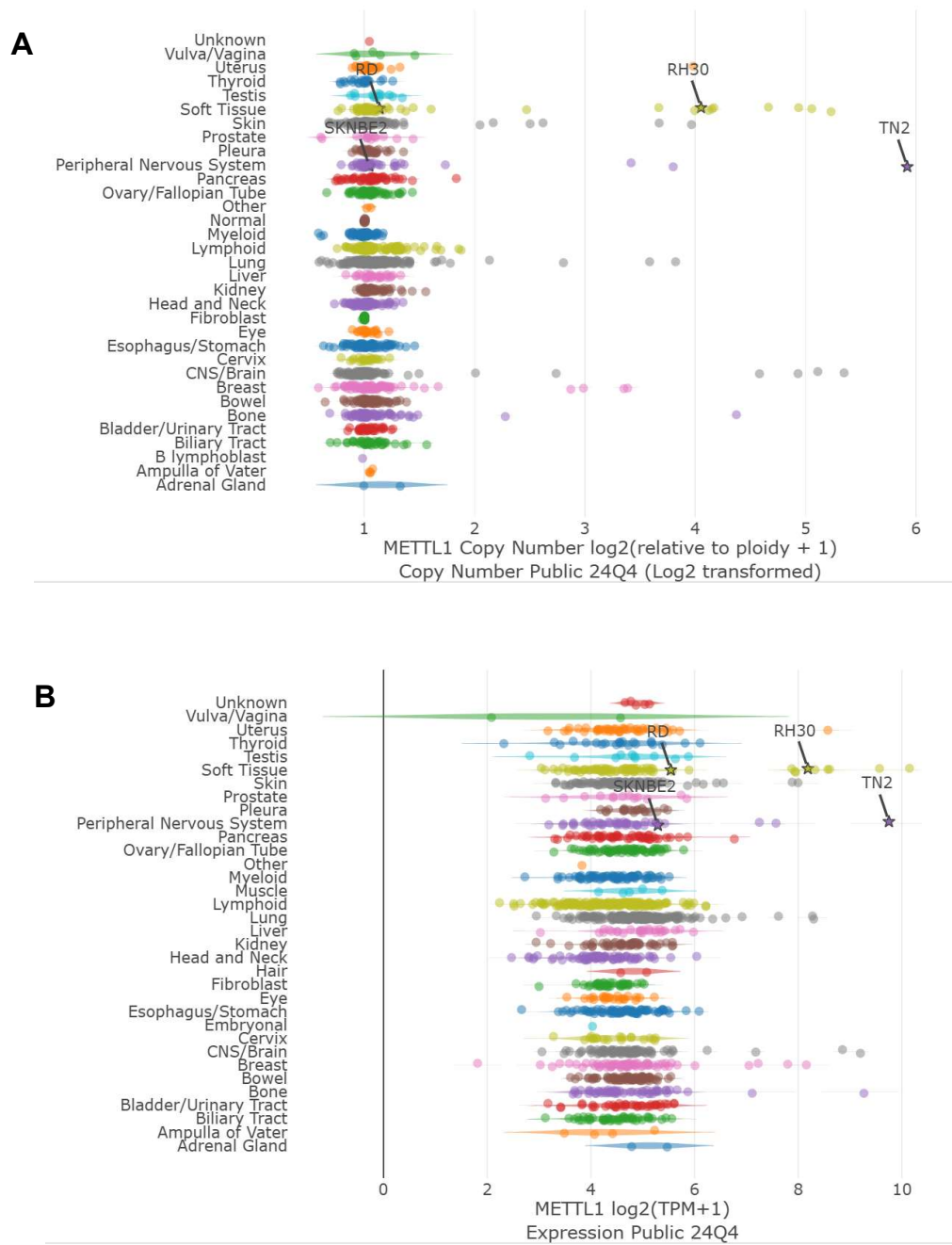

**Fig. S1. *METTL1* is amplified and overexpressed in a subset of pediatric cancer cell lines. (A-B) *METTL1* gene copy number (A) and gene expression (B) in 1178 cell lines from the CRISPR 24Q4+Score Chronos project in DepMap.**

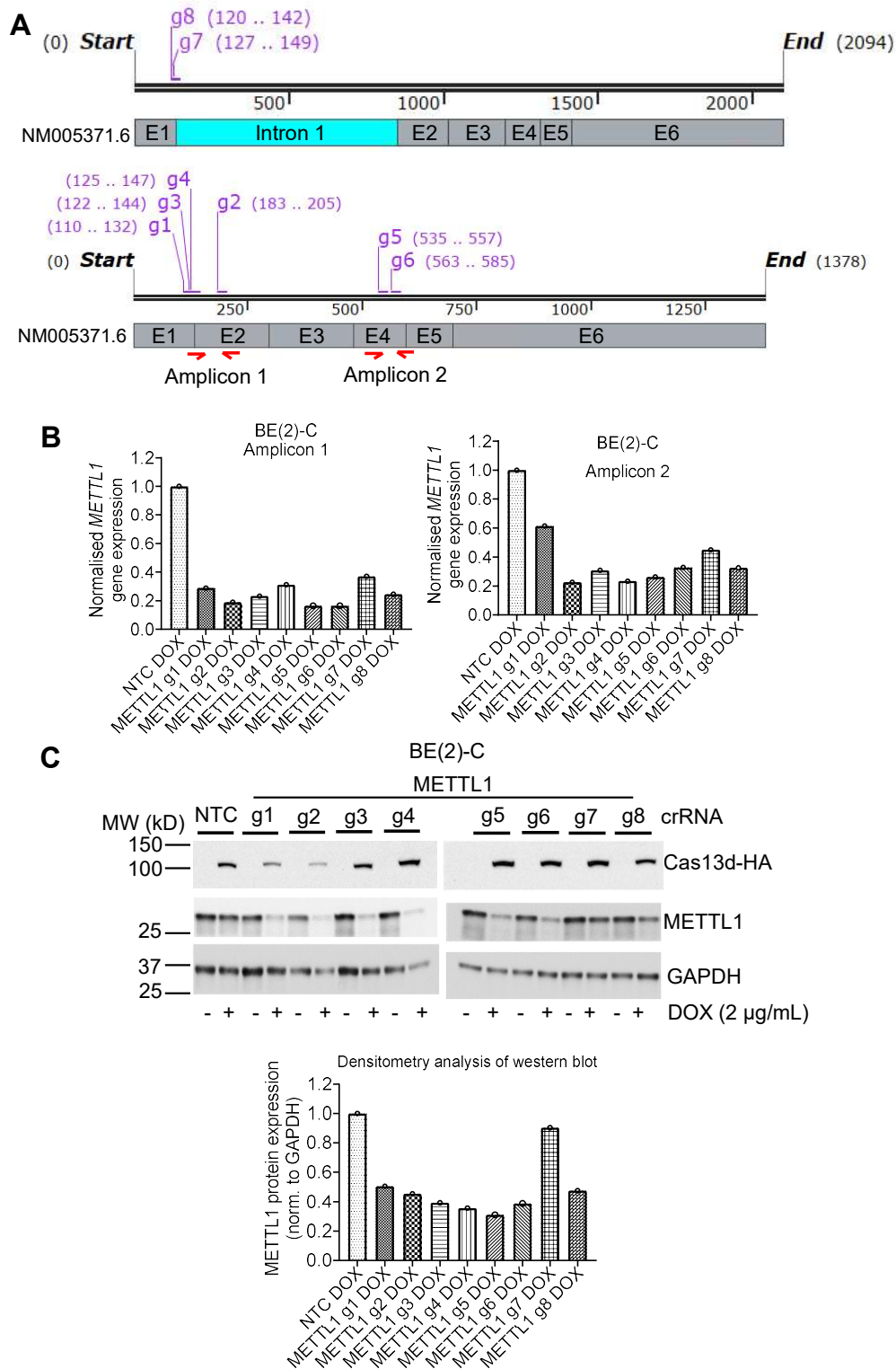

**Fig. S2. CRISPR-Cas13d targeting effectively reduces METTL1 expression.** (A) A schematic of eight METTL1 guide crRNA target regions relative to the NM005371.6 METTL1 transcript. The NM005371.6 transcript presented in the top panel includes intron 1 to depict the g7 and g8 targets in METTL1 pre-mRNA. Numbers in brackets next to the guide crRNAs indicate the positions of the guide-binding sites. Amplicons 1 and 2 are PCR products from primer pairs 1 and 2, respectively, and are marked by red arrows. The figure is not drawn to scale. E, exon. (B-C) BE(2)-C cells expressing DOX-inducible Cas13d and a single constitutively expressed crRNA (NTC or METTL1 g1, 2, 3, 4, 5, 6, 7 or 8) were treated with vehicle control or DOX for 96 hours, followed by RT-PCR (B) and western blot (C) analyses of the indicated protein expression.

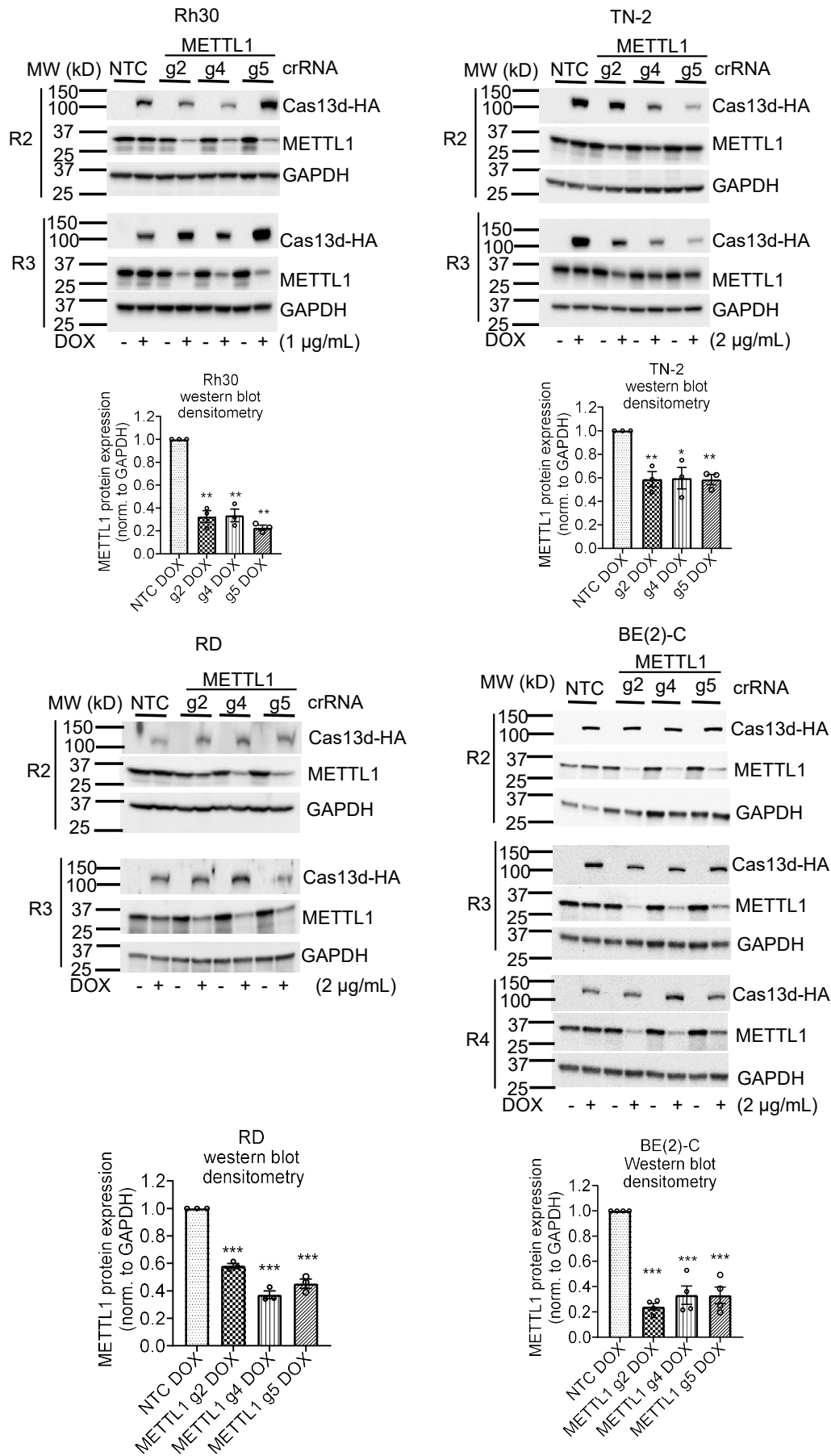

**Fig. S3. Additional western blots and densitometry analyses show that CRISPR-Cas13d targeting effectively reduces METTL1 protein expression.** The indicated cells expressing DOX-inducible Cas13d and a single constitutively expressed crRNA (NTC or METTL1 g2, g4 or g5) were treated with vehicle control or DOX for 48 (Rh30 and TN-2), 72 (RD) or 96 [BE(2)-C] hours followed by western blot and densitometry analyses of METTL1 expression. Data are presented as mean ± s.e.m. \*, \*\* & \*\*\* indicate $P < 0.05$, 0.01 & 0.001 respectively by comparing to NTC DOX treated using unpaired T-test. R2, replicate 2; R3, replicate 3; R4, replicate 4. R1 western blot data are presented in Fig. 2C.

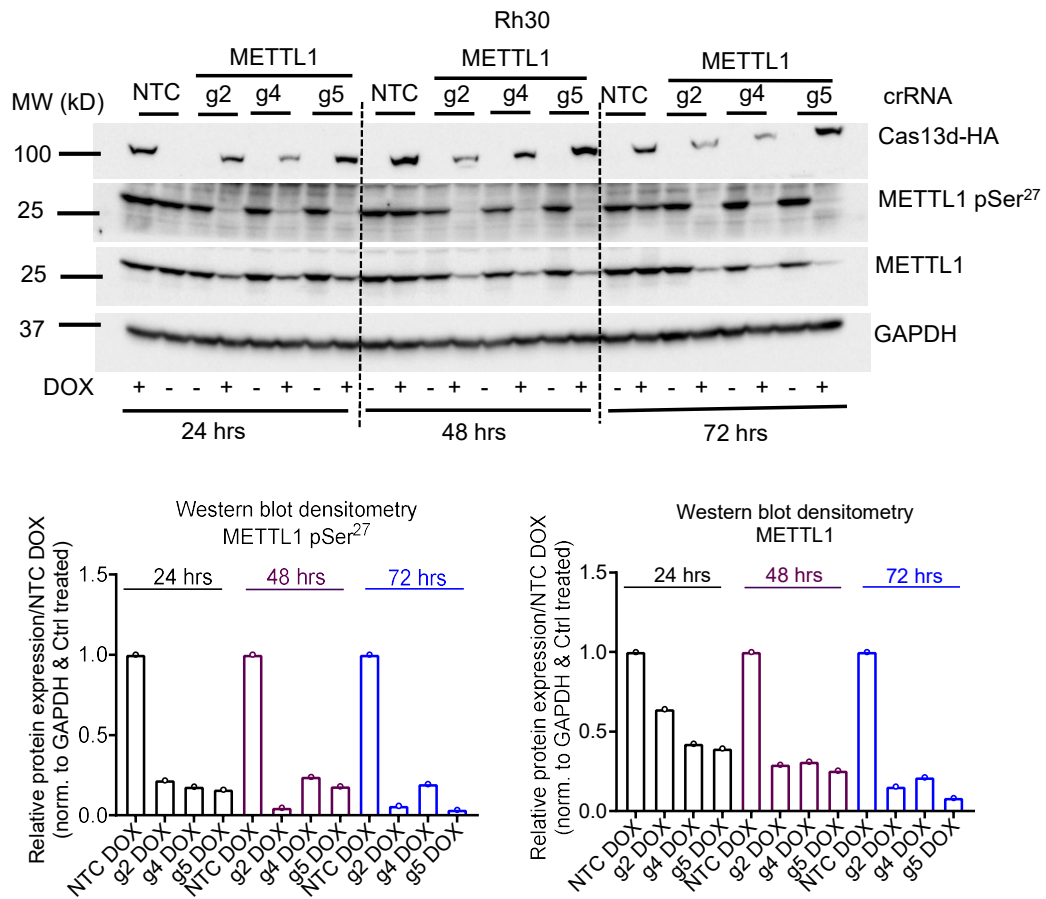

**Fig. S4. CRISPR-Cas13d targeting effectively reduces the expression of phosphorylated METTL1 protein.** Rh30 cells expressing DOX-inducible Cas13d and a single constitutively expressed crRNA (NTC or METTL1 g2, g4 or g5) were treated with vehicle control or DOX for 24, 48 or 72 hours followed by western blot and densitometry analyses of the indicated protein expression.

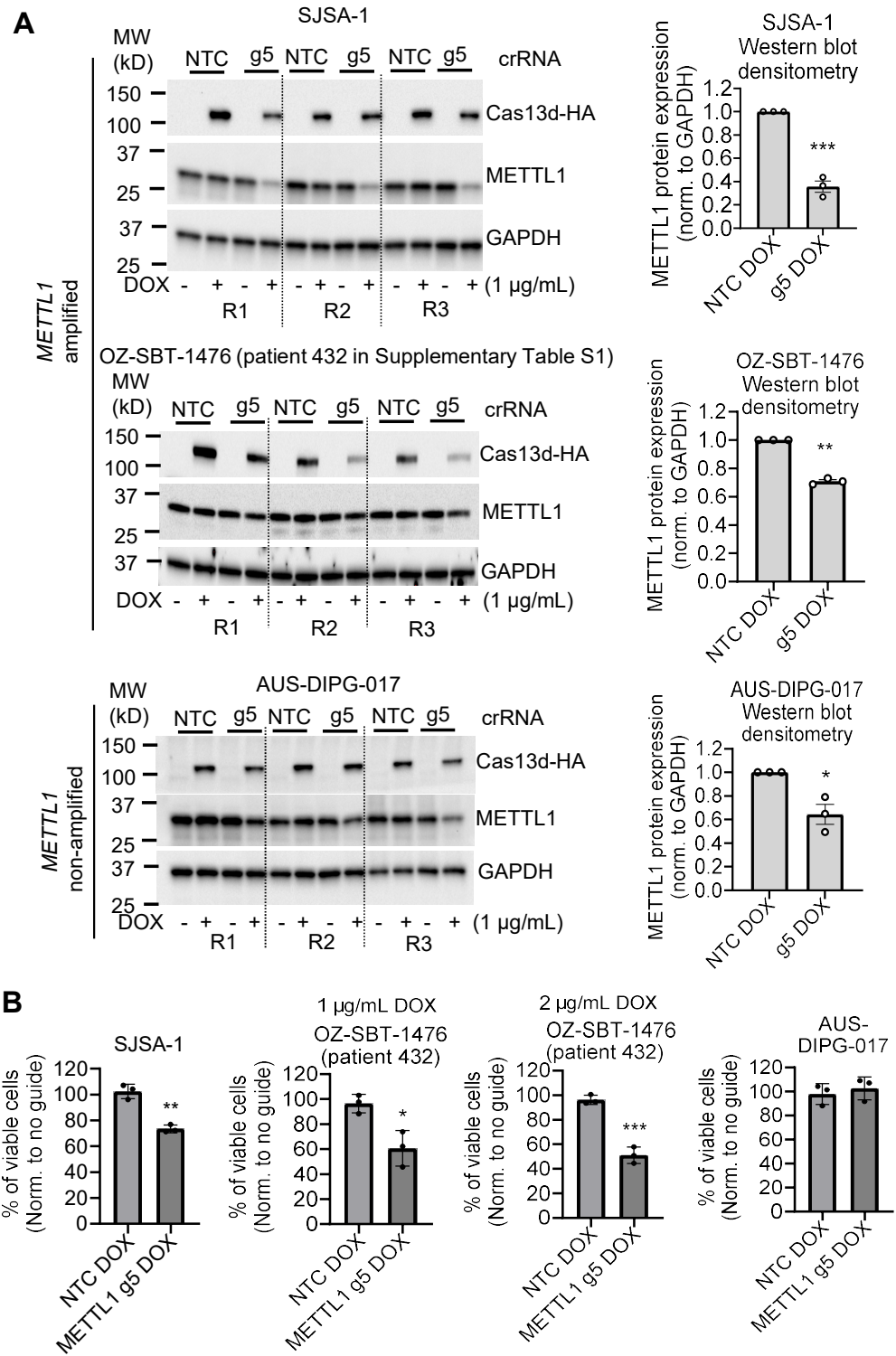

**Fig. S5. Cas13d-dependent METTL1 silencing reduces survival of *METTL1* amplified cancer cell lines. (A-B)** Cells expressing DOX-inducible Cas13d and a single constitutively expressed crRNA (NTC or METTL1 g5) were treated with vehicle control or DOX for 2 or 4 days followed by western blot analysis of METTL1 protein expression (A) or cell viability assays (B) respectively. Data are presented as mean ± s.e.m. \*, \*\* & \*\*\* indicate $P < 0.05$, 0.01 & 0.001 respectively by comparing to NTC using unpaired T-test. R1, replicate 1; R2, replicate 2; R3, replicate 3.

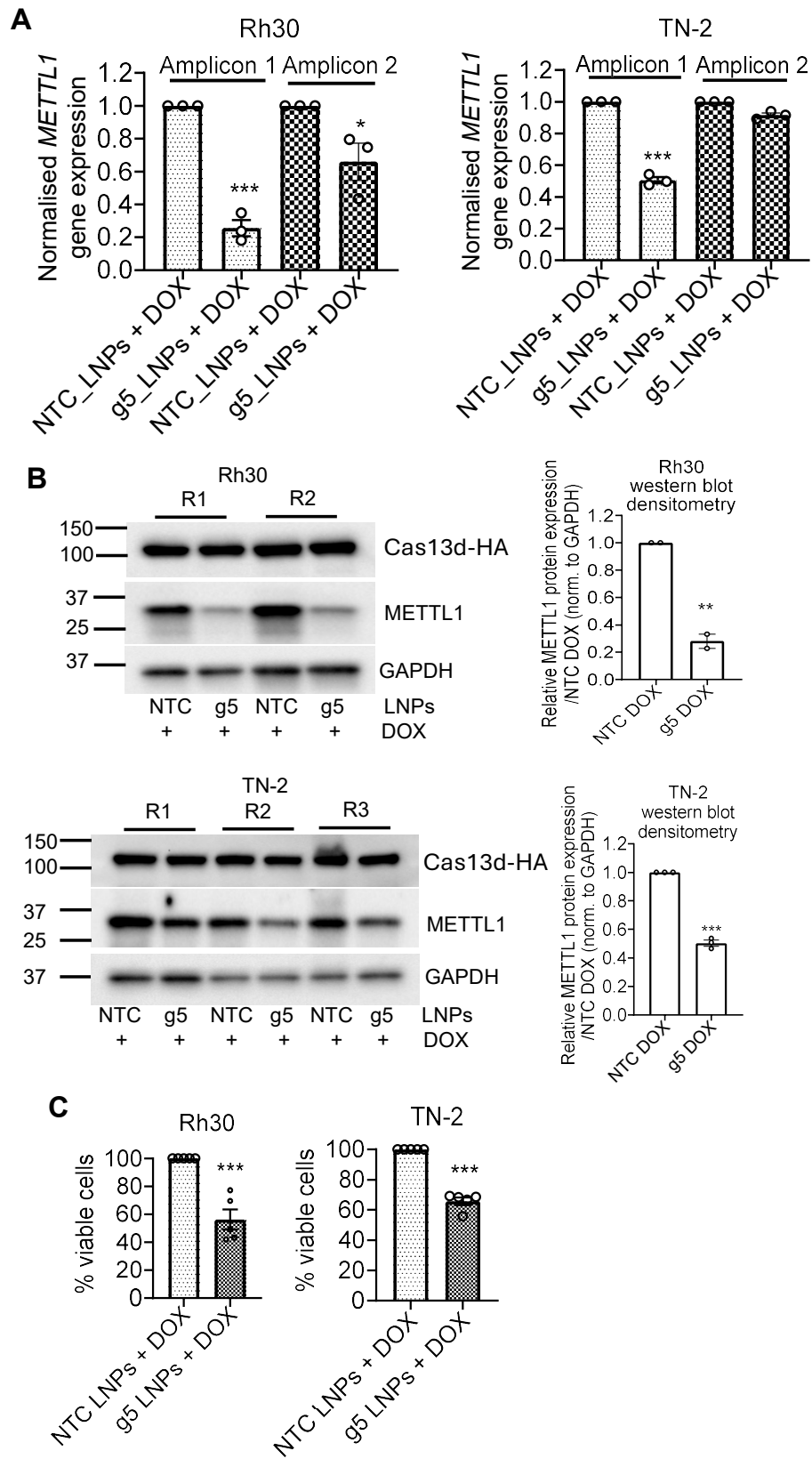

**Fig. S6. METTL1 g5 crRNA packaged in LNPs effectively reduces METTL1 expression and cell viability.** (A-C) Rh30 and TN-2 cells expressing DOX-inducible Cas13d were treated with DOX (1 μg/mL for Rh30 and 2 μg/mL for TN-2) for 12 hours, then treated with NTC or METTL1 g5 crRNA packaged in LNPs, followed by RT-PCR at 48 hours after LNP treatment (A), and western blot and densitometry (B) analyses of METTL1 expression at 48 hours after LNP treatment (B); or cell viability assays at 120 hours after LNP treatment (C). Amplicons 1 and 2 are PCR products from primer pairs 1 and 2, respectively (refer to Fig. 2A for primer target regions). Data are presented as mean ± s.e.m. \*, \*\* and \*\*\* indicate $P < 0.05$, 0.01 and 0.001, respectively, compared with DOX-treated NTC using an unpaired $t$-test. R1, replicate 1; R2, replicate 2; R3, replicate 3. Additional Rh30 western blot replicates shown in B were performed together with the puromycin incorporation assay presented in Fig. S13A.

**A** 4 independent repeats

BE(2)-C
NTC
Or METTL1 g2
Or METTL1 g4
Or METTL1 g5

Ctrl
96 hrs
DOX
96 hrs

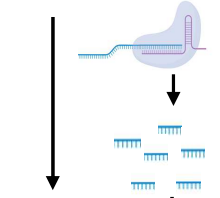

Harvest
RNA
16
samples
Harvest
RNA
16
samples

Whole
transcriptome
sequencing

**B**

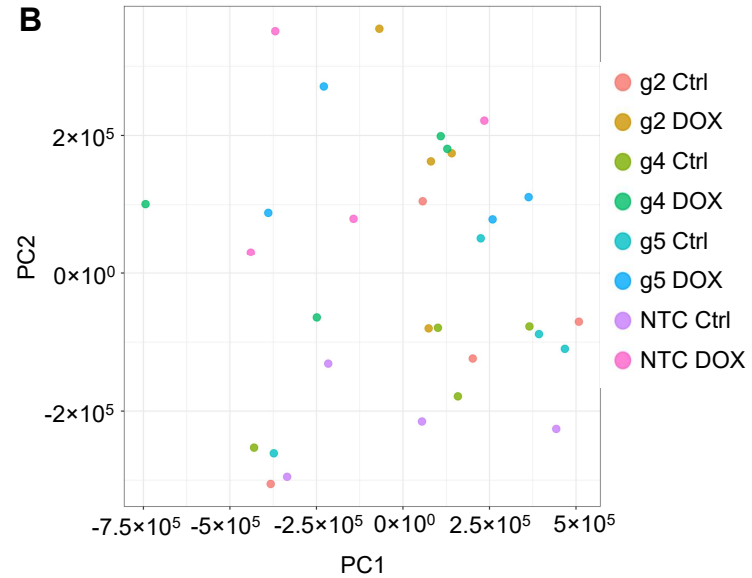

**D**

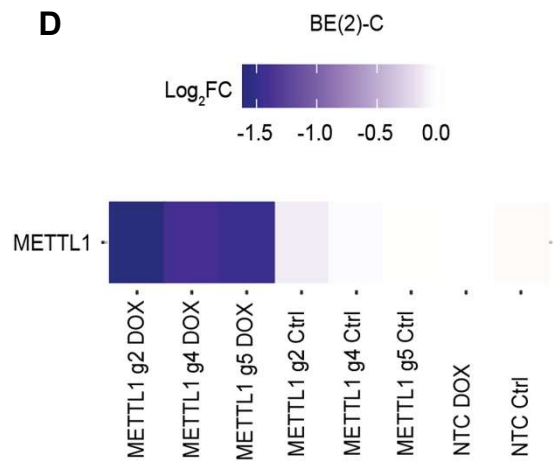

**C**

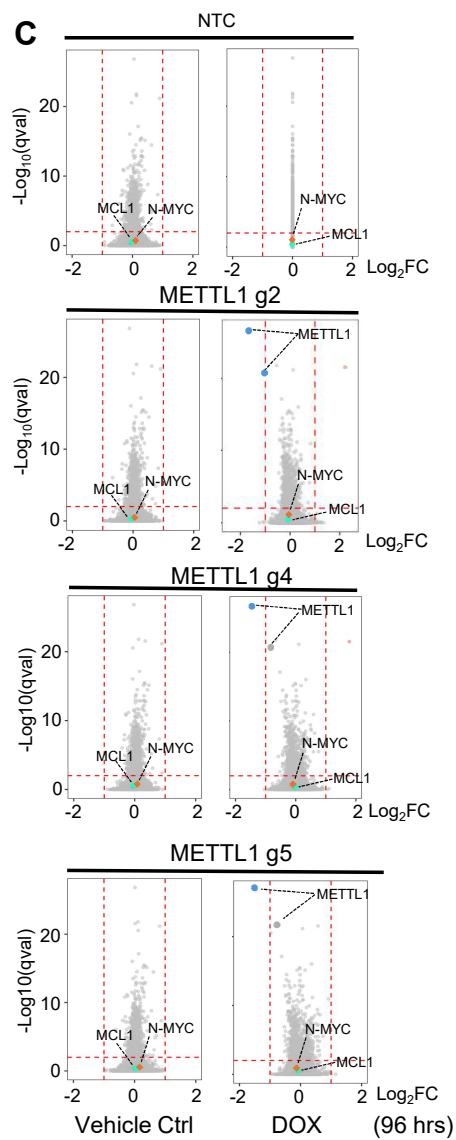

**E**

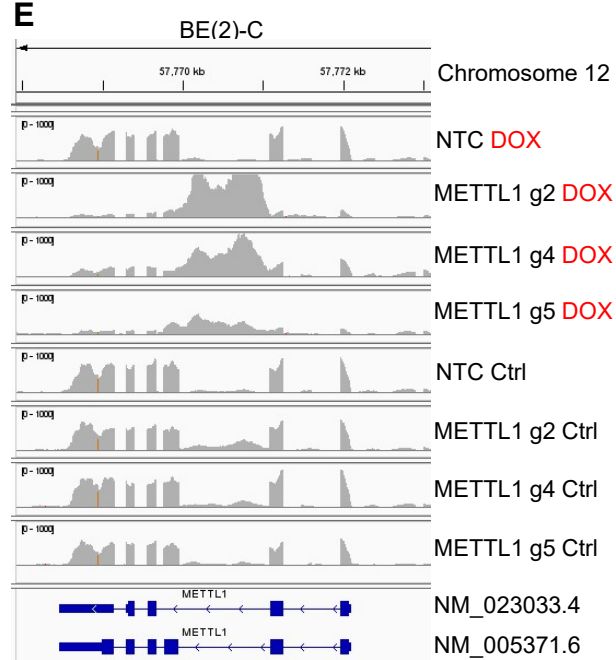

**Fig. S7. Transcriptional consequences of *METTL1* mRNA silencing are specific in BE(2)-C.** (A) A schematic of experimental design for whole transcriptome sequencing after *METTL1* silencing in BE(2)-C cells. (B) BE(2)-C cells expressing DOX-inducible Cas13d and a single constitutively expressed crRNA (NTC or *METTL1* g2, g4 or g5) were treated with vehicle control or DOX for 96 hours, followed by whole transcriptome sequencing and principal component analysis. (C) Transcriptome wide differential transcript expression (DTE) analysis showing the common transcripts up- or down-regulated in four independent experimental repeats as volcano plots. The dotted lines mark the cut-off points using log2 fold-change $<-1$ or $>1$ and $qval < 0.01$. Blue dots on the left of the x-axis and orange dots on the right of the x-axis represent the genes that are significantly down or upregulated respectively after *METTL1* silencing. (D) The common significantly differentially expressed canonical transcripts in the four independent experimental repeats are presented as a heatmap. (E) A diagram showing the well-supported *METTL1* transcripts (GENCODE release 48 Primary Set) after *METTL1* silencing in Integrative Genomics Viewer.

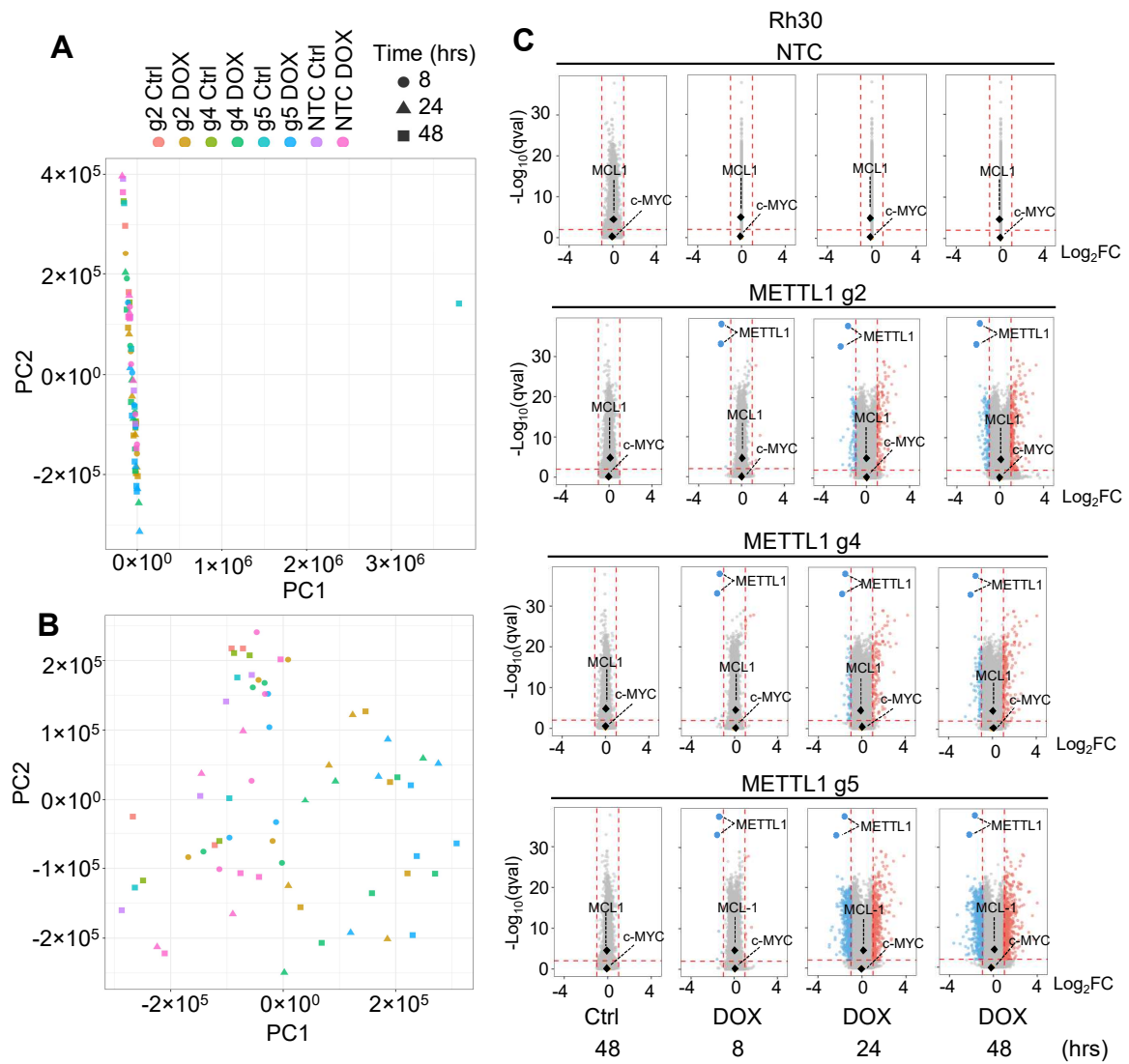

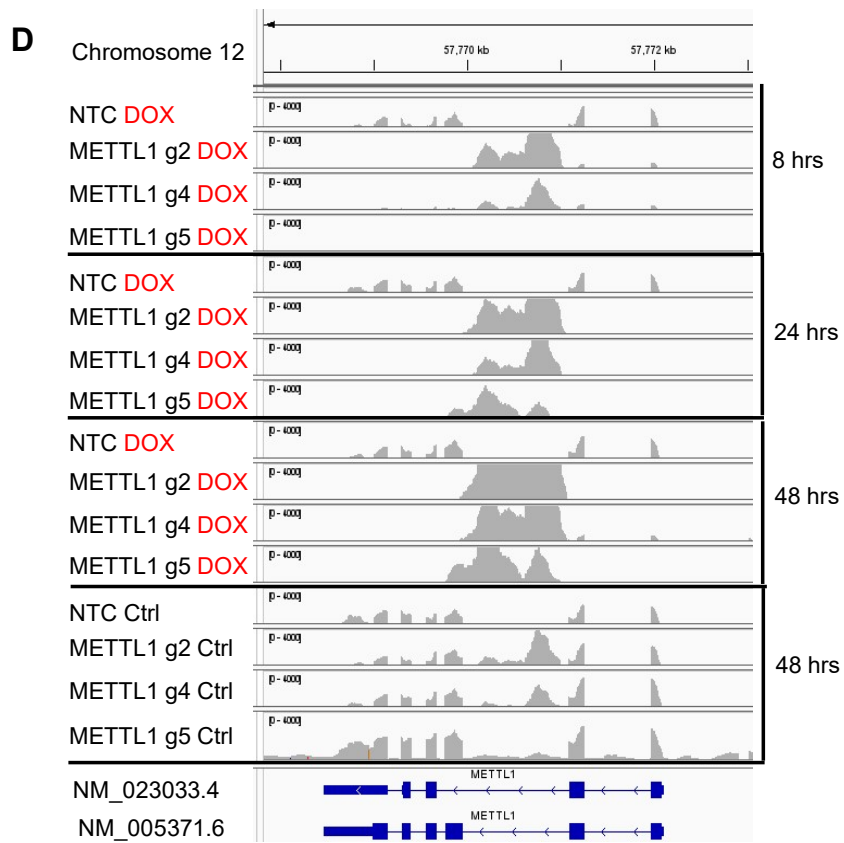

**Fig. S8. Transcriptional consequences of *METTL1* mRNA silencing are specific in Rh30.** (A-B) Rh30 cells expressing DOX-inducible Cas13d and a single constitutively expressed crRNA (NTC or *METTL1* guide 2, g4 or g5) were treated with vehicle control or DOX for 8, 24 and 48 hours followed by whole transcriptome sequencing and principal component analysis (A) and reanalysis by removing an outlier, g5 control treated for 48 hours replicate 3 (B). (C) Transcriptome wide differential transcript expression (DTE) analysis showing the common transcripts up or down regulated in four independent experimental repeats as volcano plots (g5 is also shown in Fig. 5B). The vertical dotted lines mark the cut-off points using $\log_2$ fold-change $<-1$ or $>1$ and the horizontal dotted line marks the cut-off point using $qval < 0.01$. Blue dots on the left of the x-axis and orange dots on the right of the x-axis represent the transcripts that are significantly down or upregulated respectively after *METTL1* silencing. (D) A diagram showing the well-supported *METTL1* transcripts (GENCODE release 48 Primary Set) after *METTL1* silencing in Integrative Genomics Viewer.

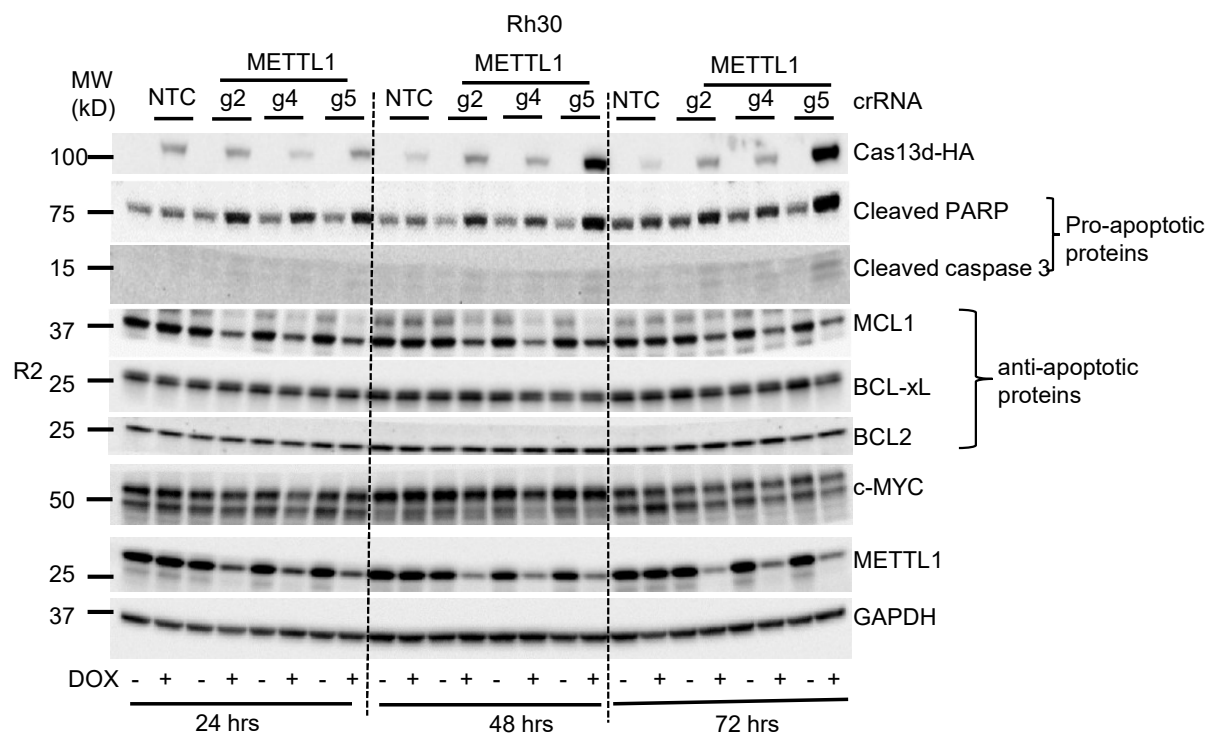

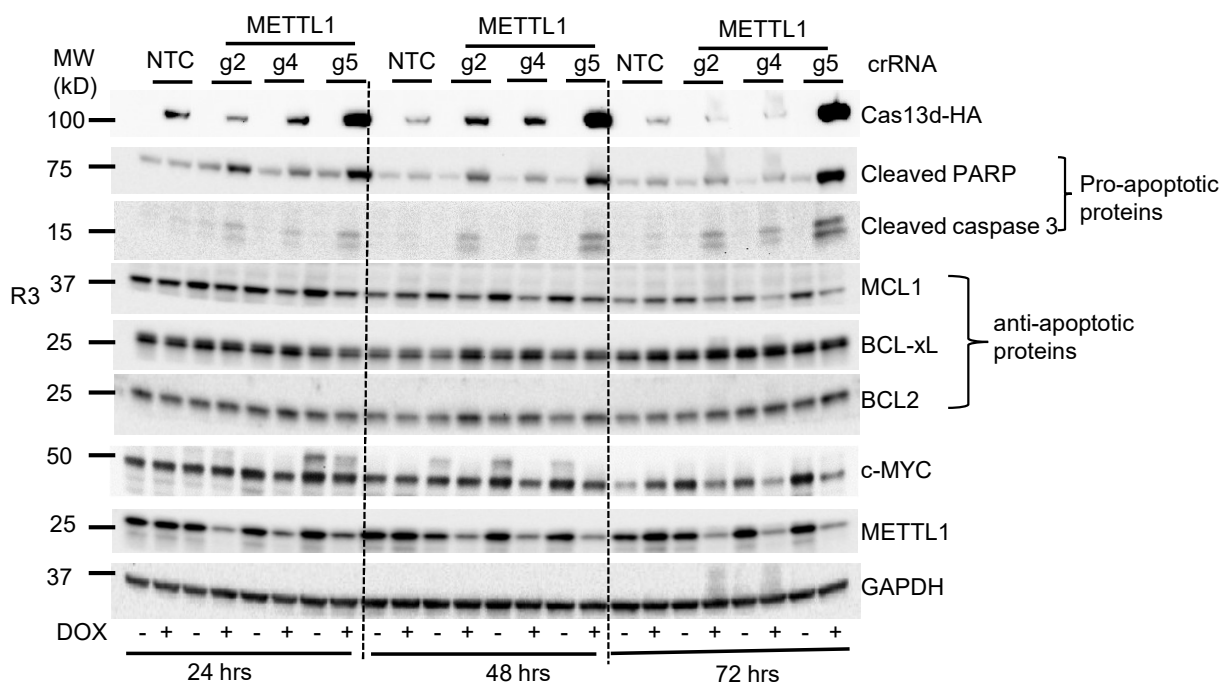

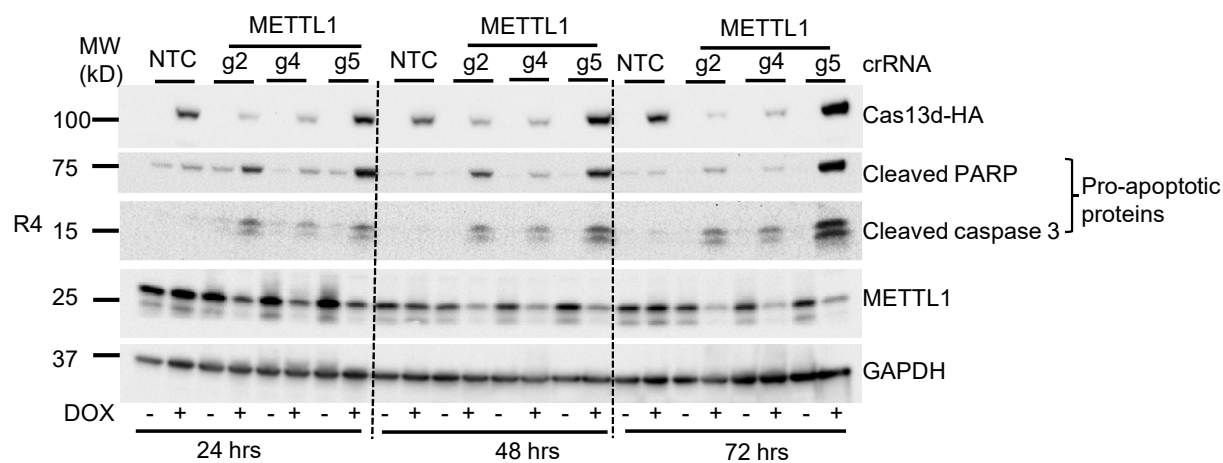

**Fig. S9. Additional western blot analyses show downregulation of METTL1 is associated with reduced MCL1 and c-MYC protein expression in Rh30.** Rh30 cells expressing DOX-inducible Cas13d and a single constitutively expressed crRNA (NTC or METTL1 g2, g4 or g5) were treated with vehicle control or DOX for 24, 48 and 72 hours, followed by western blot analysis of the indicated protein expression. R2, replicate 2; R3, replicate 3; R4, replicate 4. R1 western blot data are presented in Fig. 6A.

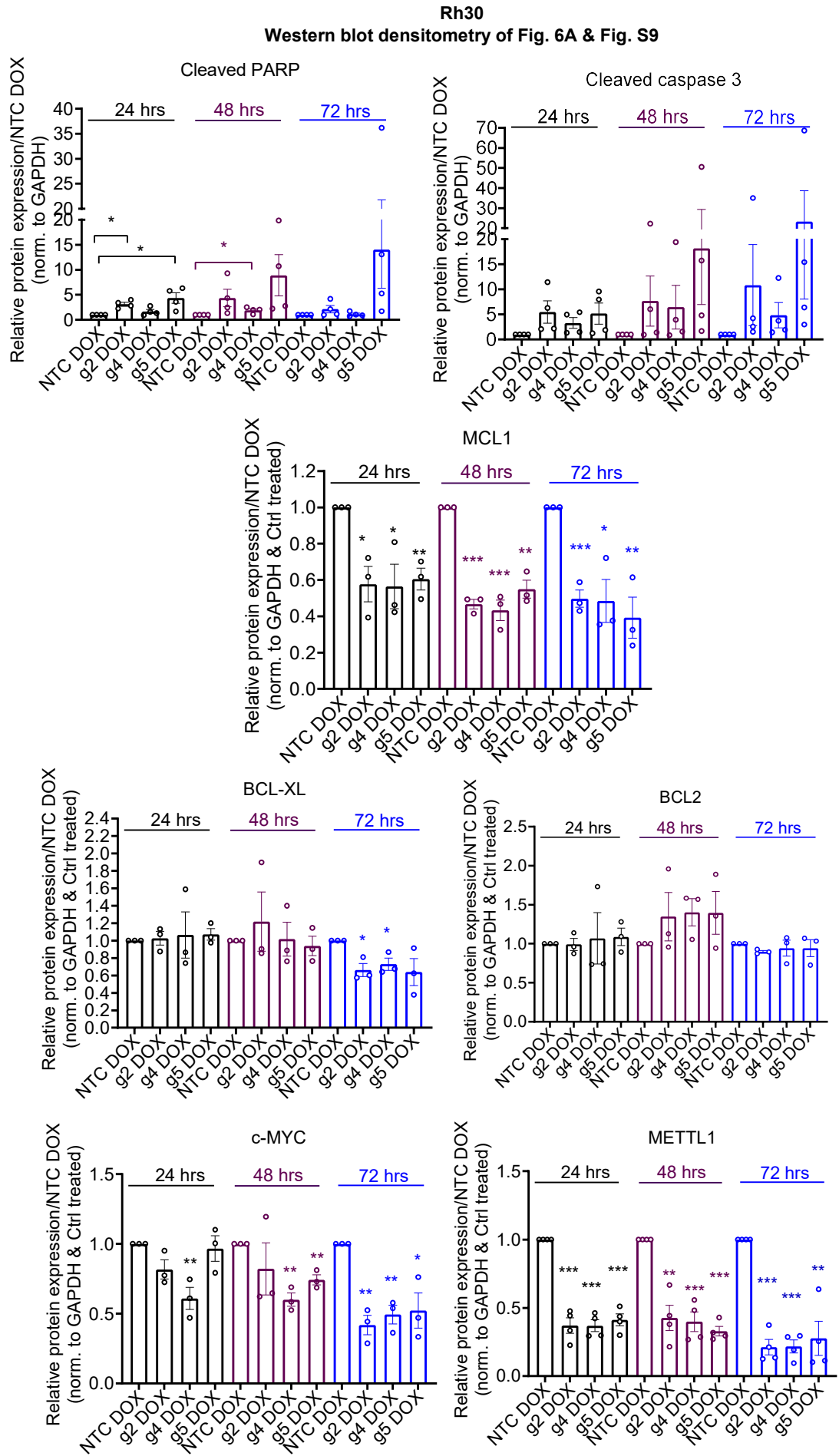

**Fig. S10. Densitometry analyses of western blots show downregulation of METTL1 is associated with reduced MCL1 and c-MYC protein expression in Rh30.** Densitometry analyses of western blot data presented in Fig. 6A and Fig. S9. Data are presented as mean ± s.e.m. \*, \*\* & \*\*\* indicate $P < 0.05$, 0.01 & 0.001 respectively by comparing to NTC DOX treated using unpaired T-test.

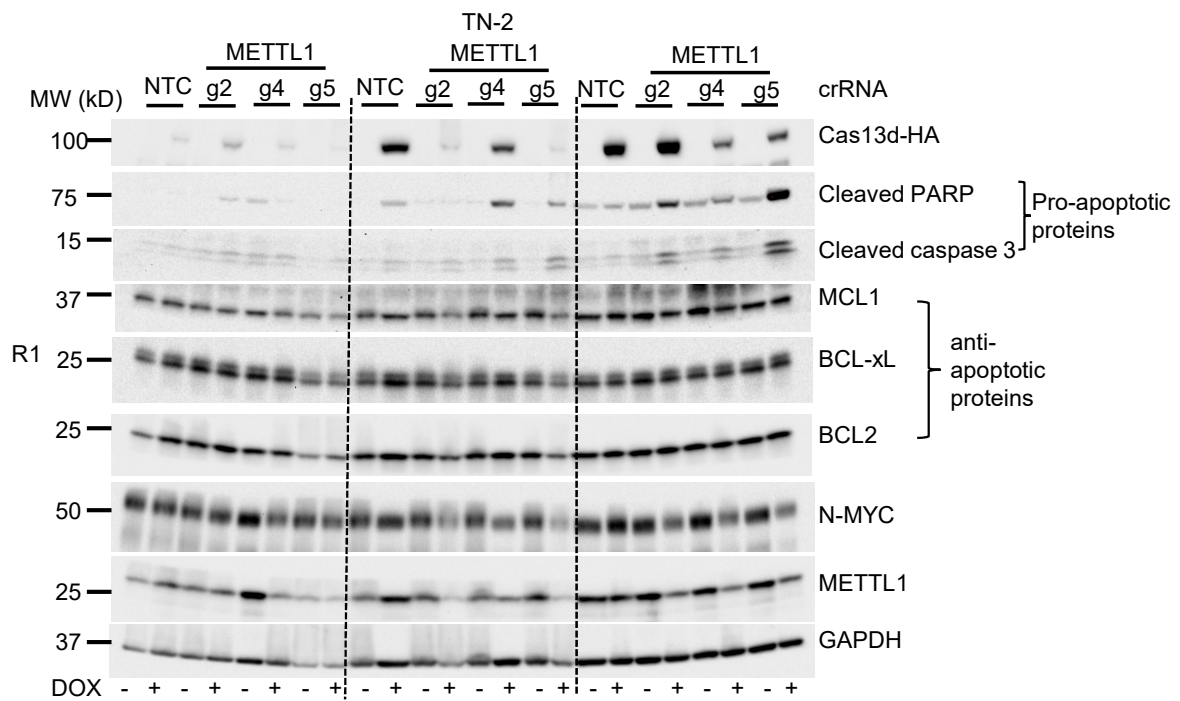

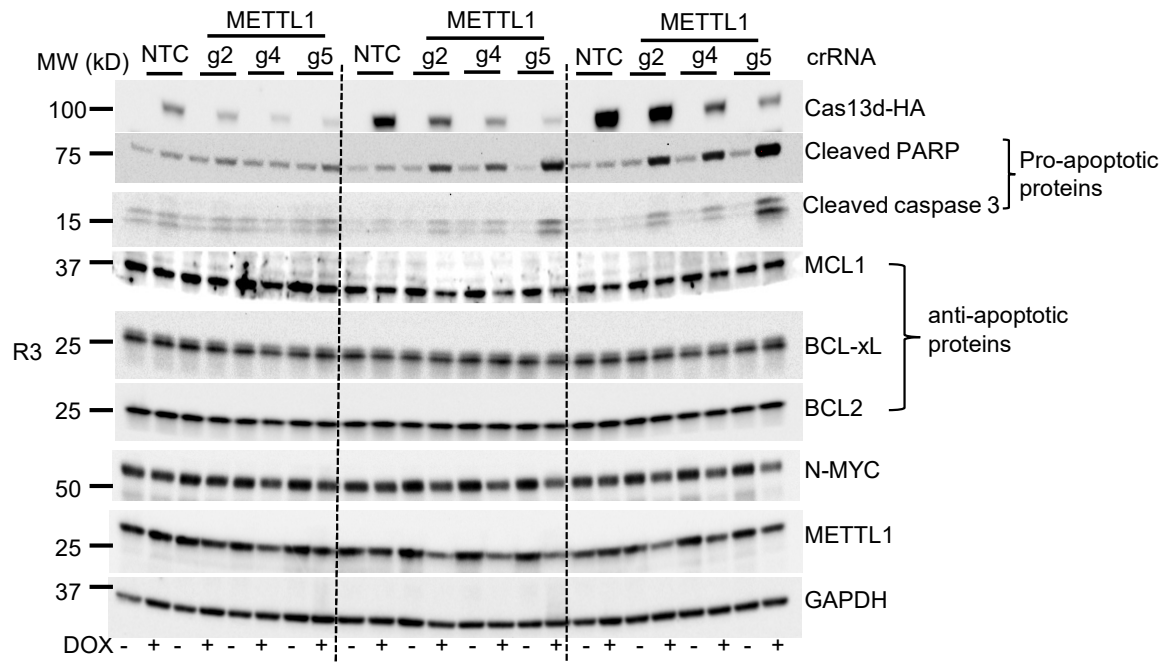

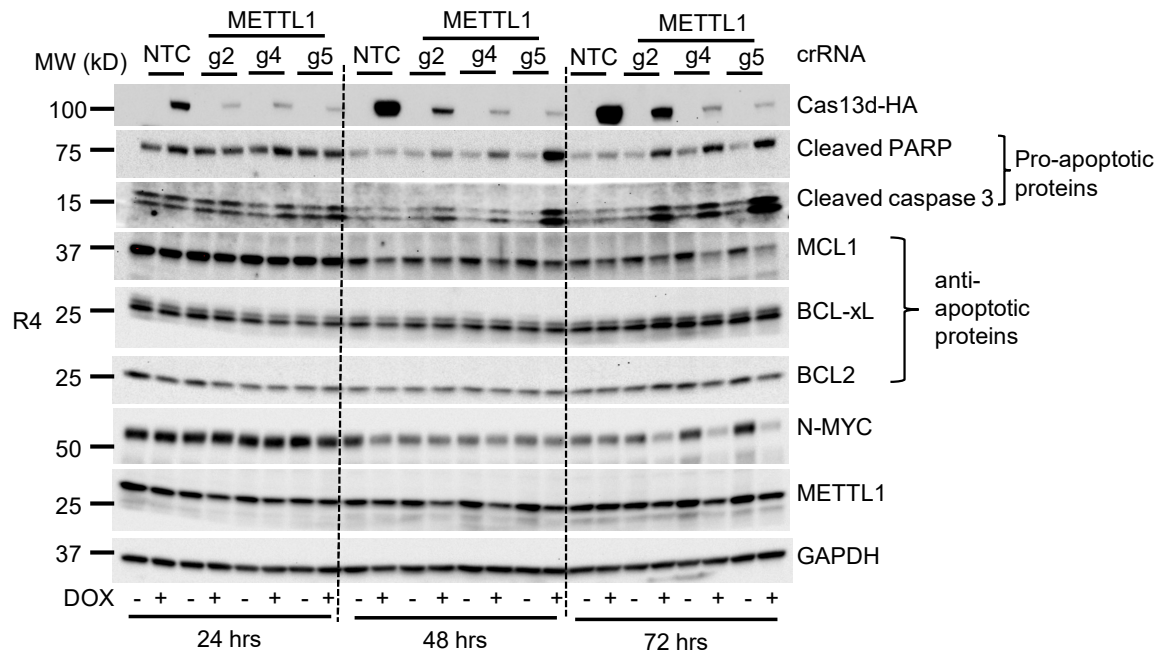

**Fig. S11. Additional western blot analyses show downregulation of METTL1 is associated with reduced N-MYC protein expression in TN-2.** TN-2 cells expressing DOX-inducible Cas13d and a single constitutively expressed crRNA (NTC or METTL1 g2, g4 or g5) were treated with vehicle control or DOX for 24, 48 and 72 hours, followed by western blot analysis of the indicated protein expression. R1, replicate 1; R3, replicate 3; R4, replicate 4. R2 western blot data are presented in Fig. 6B.

**TN-2**
**Western blot densitometry of Fig. 6B & Fig. S11**

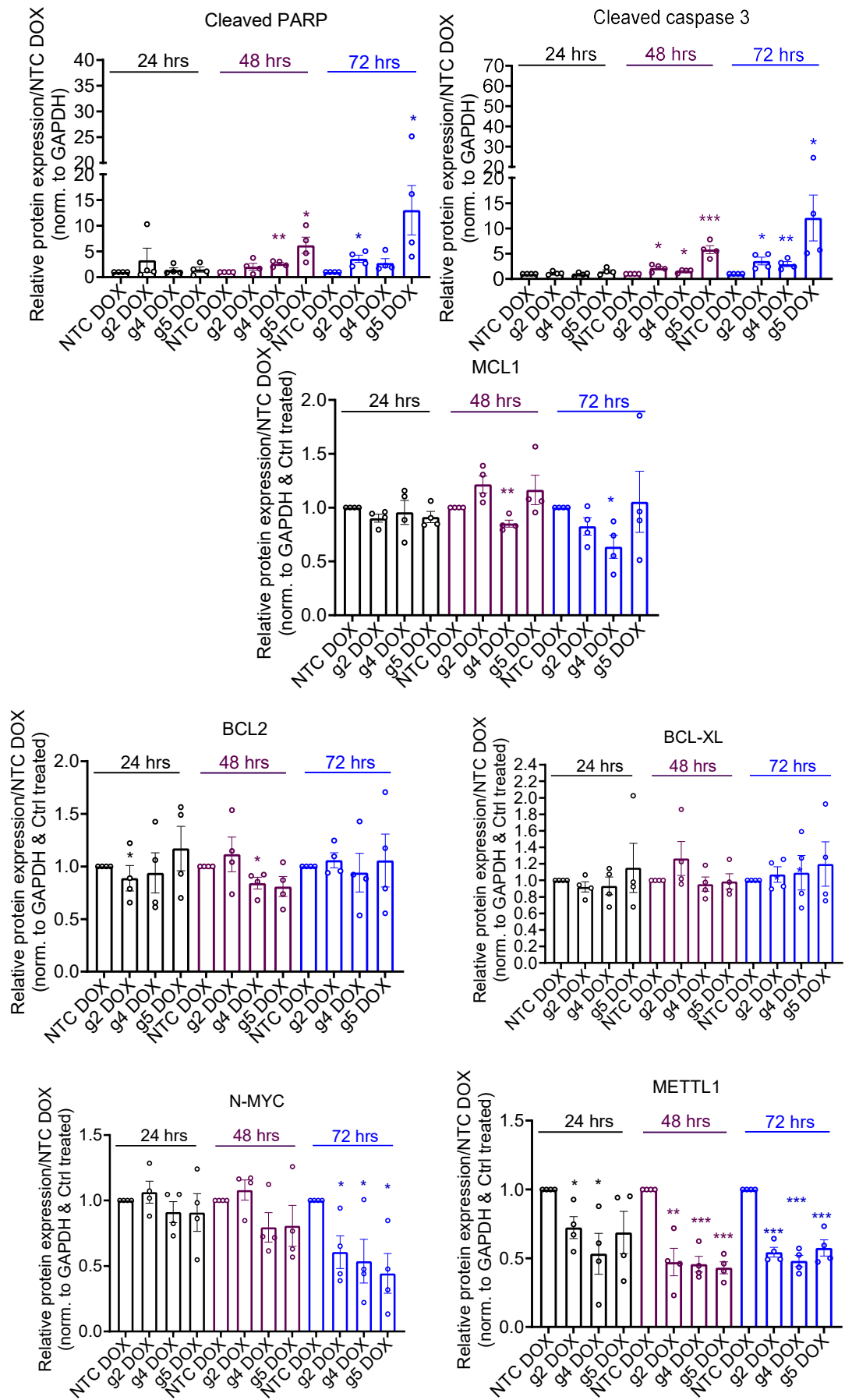

**Fig. S12. Densitometry analyses of western blots show downregulation of METTL1 is associated with reduced N-MYC protein expression in TN-2.** Densitometry analyses of western blot data presented in Fig. 6B and Fig. S11. Data are presented as mean ± s.e.m. \*, \*\* & \*\*\* indicate $P < 0.05$, $0.01$ & $0.001$ respectively by comparing to NTC DOX treated using unpaired T-test.

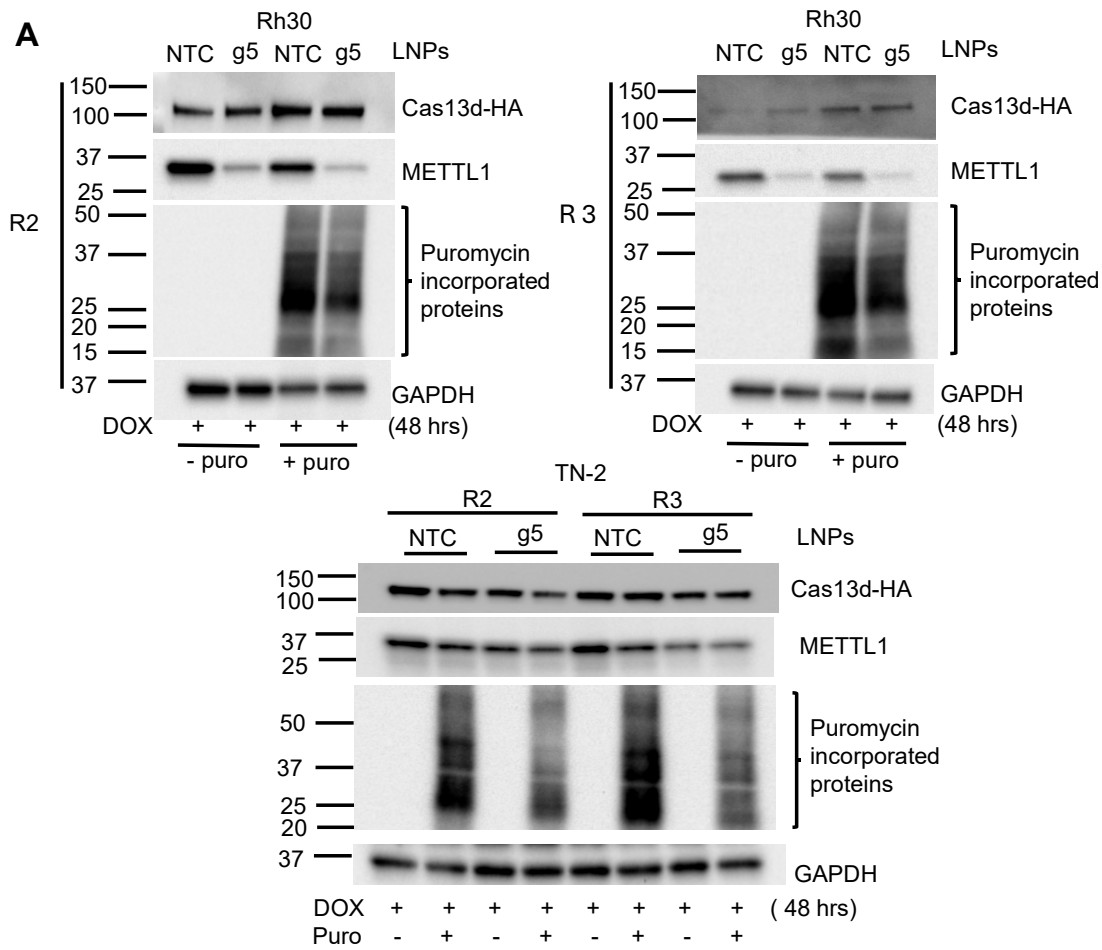

**Western blot densitometry of Fig. 7A & Fig. S13A**

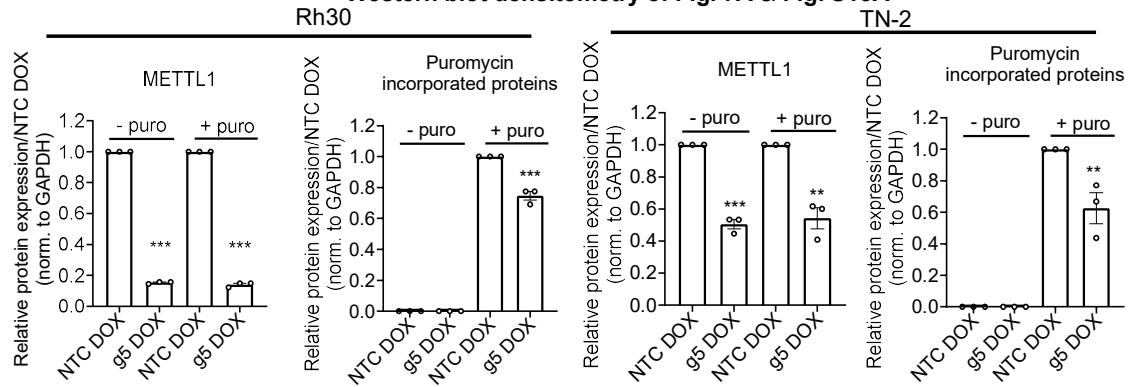

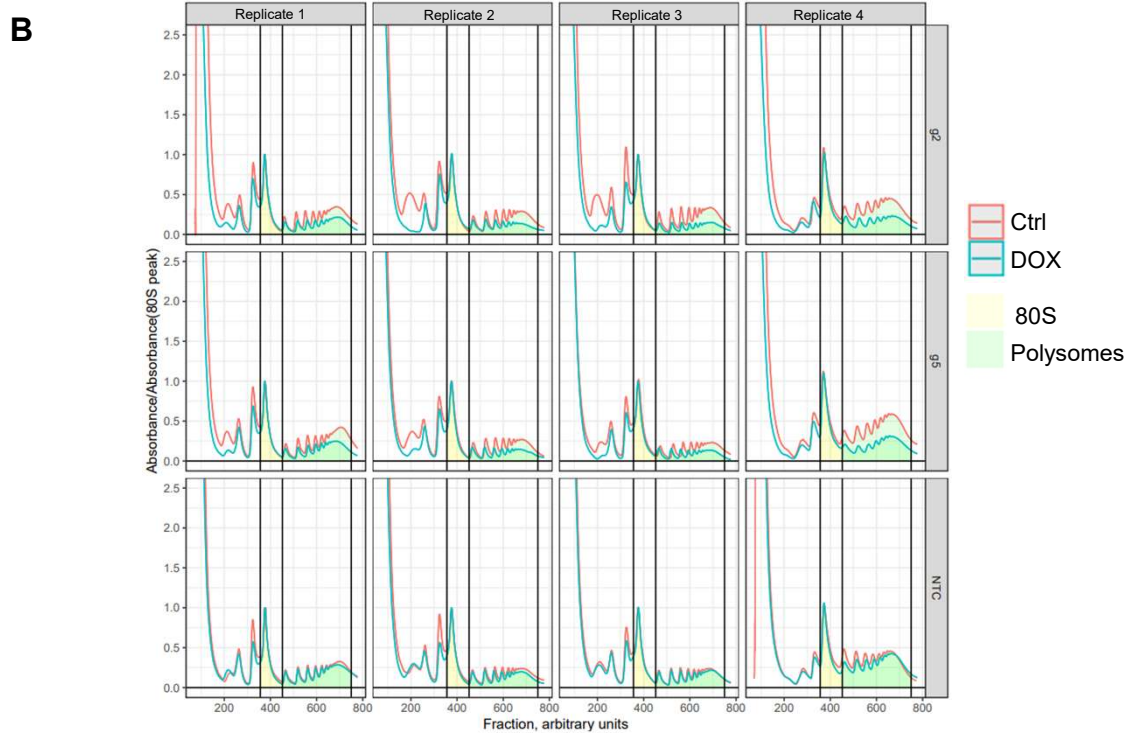

**Fig. S13. Downregulation of METTL1 is associated with reduced global protein translation.** (A) Rh30 and TN-2 cells expressing DOX-inducible Cas13d were treated with control or DOX for 12 hours, followed by the addition of NTC or METTL1 g5 crRNA packaged in LNPs. Forty-eight hours post LNPs treatment, the cells were pulsed with puromycin followed by western blot and densitometry analyses of the indicated protein expression. R2, replicate 2; R3, replicate 3. R1 western blot data are presented in Fig. 7A. (B) Rh30 cells expressing DOX-inducible Cas13d and a single constitutively expressed crRNA (NTC or g2 or g5) were treated with vehicle or DOX for 48 hours, followed by treatment with cycloheximide and polysome fractionation. Replicate 4 is also presented in Fig. 7B. Data are presented as mean ± s.e.m. \*\* & \*\*\* indicate $P < 0.01$ & $0.001$ respectively by comparing to NTC DOX treated using unpaired T-test.

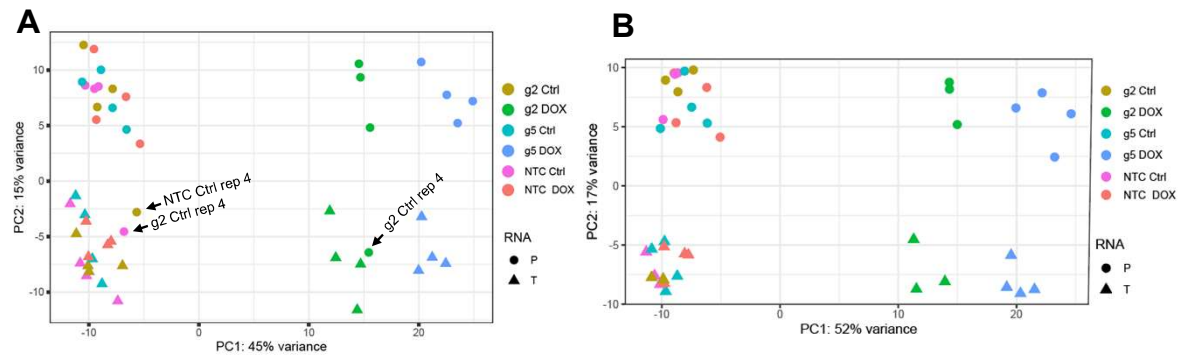

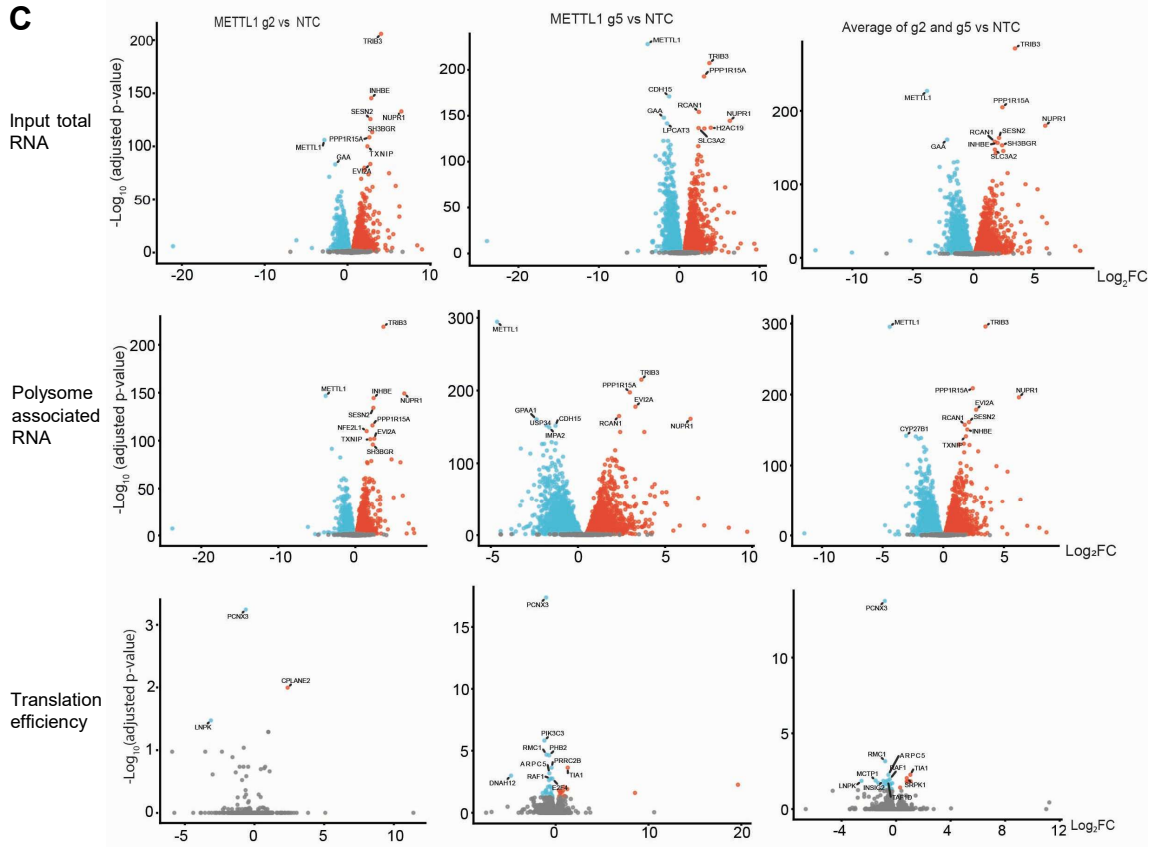

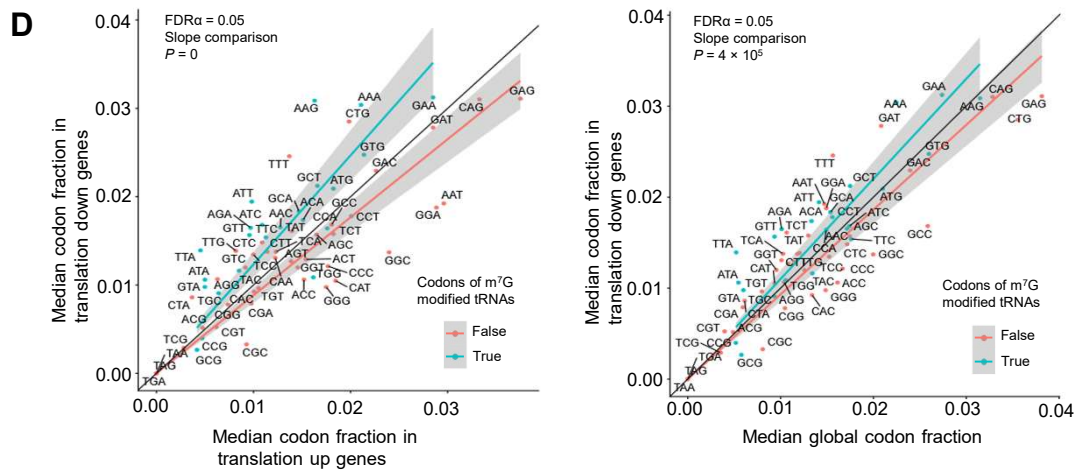

**Fig. S14. Translational consequences of *METTL1* mRNA silencing in Rh30. (A–D)** Rh30 cells expressing DOX-inducible Cas13d and a single constitutively expressed crRNA (NTC or *METTL1* g2 or g5) were treated with vehicle control or 1.25 µg/mL DOX for 48 hours, followed by polysome fractionation and RNA sequencing. (A) Principal component analysis of all samples. P, polysomal RNA; T, total RNA. (B) Principal component analysis after removal of three outliers: NTC and g2 treated with vehicle control, and g2 treated with DOX, all from polysomal RNA in replicate 4. (C) Transcriptome-wide differential gene expression analysis, shown as volcano plots. Upper panel, total RNA; middle panel, polysome-associated RNA; lower panel, translation efficiency. Each point represents a gene: blue, significantly downregulated; red, significantly upregulated; grey, not significant. Significant genes are those changed in at least three independent experimental repeats. (D) Median codon usage after *METTL1* silencing. Left panel, 20 translationally downregulated genes versus 6 translationally upregulated genes; right panel, the same 20 downregulated genes versus the global codon fraction. Each point represents a single codon, and in each panel the diagonal black line indicates equal codon usage between the two groups compared. Blue and red lines show linear regressions for codons decoded by m<sup>7</sup>G-modified and non-m<sup>7</sup>G-modified tRNAs, respectively. Grey shaded areas correspond to the 95% confidence interval for the predicted regression.

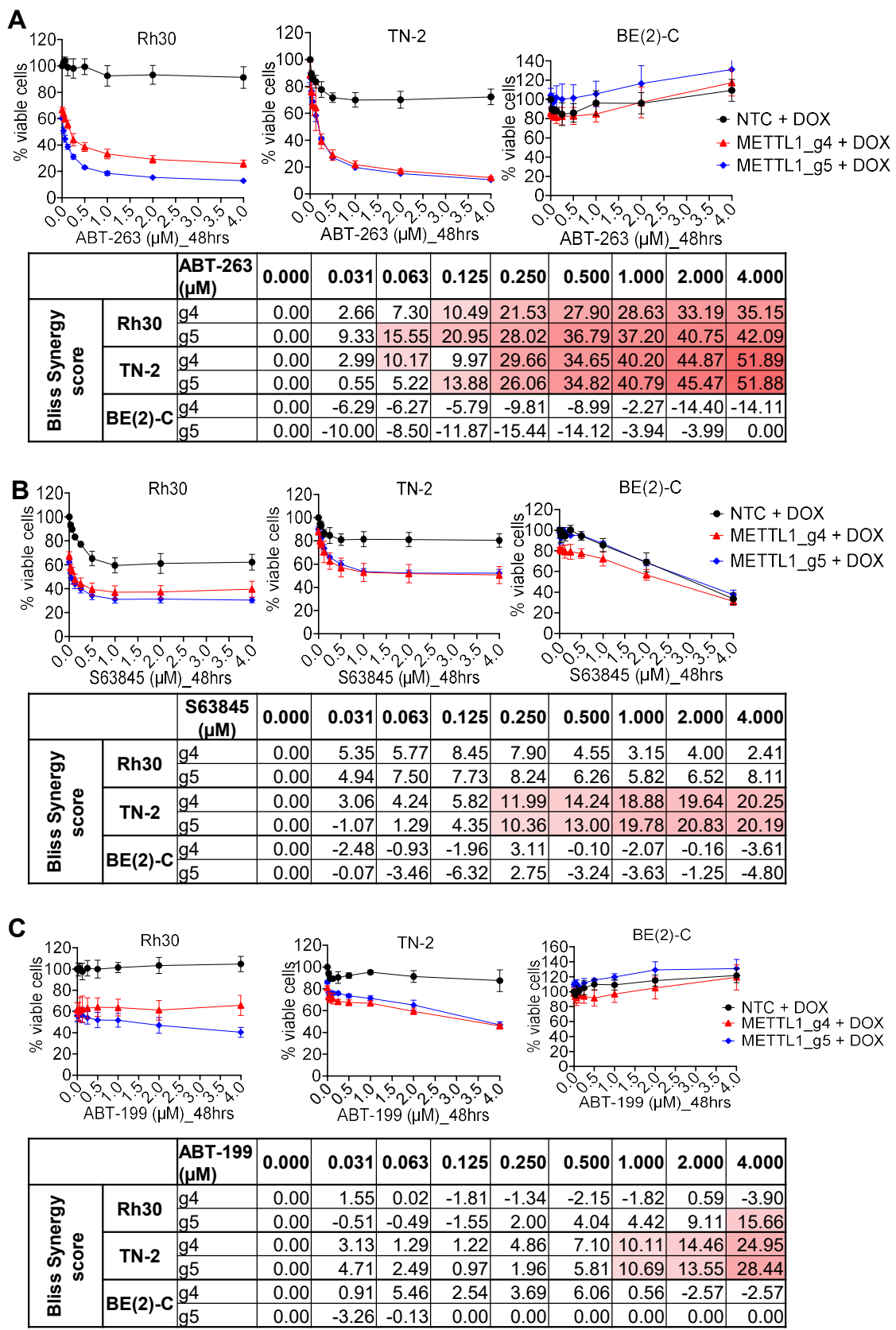

**Fig. S15. METTL1 inhibition synergizes with a BCL2 family protein inhibitor. (A-C)** Cells expressing DOX-inducible Cas13d and a single constitutively expressed crRNA (NTC or METTL1 g4 or g5) were treated with vehicle control or DOX [0.5 μg/mL for Rh30, 1 μg/mL for BE(2)-C and TN-2] and ABT-263, S63845 or ABT-199 at the indicated concentrations for 48 hours, followed by cell viability assays. Bliss synergy scores > 10 and 15 indicate synergy and strong synergy respectively.
